# Autophagy-dependent TOR reactivation drives fungal growth in living host rice cells

**DOI:** 10.1101/2021.10.20.465203

**Authors:** Gang Li, Ziwen Gong, Nawaraj Dulal, Raquel O. Rocha, Richard A. Wilson

## Abstract

Eukaryotic filamentous plant pathogens with biotrophic growth stages like the devastating hemibiotrophic rice blast fungus *Magnaporthe oryzae* grow for extended periods in living host plant cells without eliciting defense responses. *M. oryzae* elaborates invasive hyphae (IH) that grow in and between living rice cells while separated from host cytoplasm by plant-derived membrane interfaces. However, although critical to the plant infection process, the molecular mechanisms and metabolic strategies underpinning this intracellular growth phase are poorly understood. Eukaryotic cell growth depends on activated target-of-rapamycin (TOR) kinase signaling, which inhibits autophagy. Here, using live-cell imaging coupled with plate growth tests and RNAseq, proteomic, quantitative phosphoproteomics and metabolic approaches, we show how cycles of autophagy in IH modulate TOR reactivation via α-ketoglutarate to sustain biotrophic growth and maintain biotrophic interfacial membrane integrity in host rice cells. Deleting the *M. oryzae* serine-threonine protein kinase Rim15-encoding gene attenuated biotrophic growth, disrupted interfacial membrane integrity and abolished the *in planta* autophagic cycling we observe here for the first time in wild type. *Δrim15* was also impaired for glutaminolysis and depleted for α-ketoglutarate. α-ketoglutarate treatment of *Δrim15-*infected leaf sheaths remediated *Δrim15* biotrophic growth. In WT, α-ketoglutarate treatment suppressed autophagy. α-ketoglutarate signaling is amino acid prototrophy- and GS-GOGAT cycle-dependent. We conclude that, following initial IH elaboration, cycles of Rim15- dependent autophagic flux liberate α-ketoglutarate – via the GS-GOGAT cycle – as an amino acid-sufficiency signal to trigger TOR reactivation and promote fungal biotrophic growth in nutrient-restricted host rice cells.

## Introduction

*Magnaporthe oryzae* (syn *Pyricularia oryzae*) (Wilson, 2021) causes blast, the most devastating disease of cultivated rice (Wilson and Talbot, 2009; Fernandez and Orth, 2018). *M. oryzae* is a hemibiotroph, an important class of eukaryotic microbial plant pathogen that, following host invasion, grows undetected for an extended period in living host plant cells before switching to necrotrophy, when host cells die and disease symptoms develop. In many pathosystems, biotrophic growth involves the elaboration of feeding structures (haustorium) or intracellular invasive hyphae (IH) that are surrounded by host plant-derived membranes to form interfacial zones for nutrient acquisition and the deployment of plant innate immunity-suppressing effectors (Yi and Valent, 2013). The *M. oryzae* biotrophic interface comprises IH wrapped in the host-derived extra-invasive hyphal membrane (EIHM), forming an interfacial compartment into which apoplastic effectors like Bas4 can be deployed (Khang et al. 2010; Giraldo et al. 2013). *M. oryzae* IH also associate with a focal plant lipid-rich structure called the biotrophic interfacial complex (BIC). A single BIC forms in the first infected rice cell following penetration at around 32 hr post inoculation (hpi) and at the tips of IH as they spread into neighbouring cells at around 44 hpi. BICs form outside IH and accumulate secreted cytoplasmic effectors such as Pwl2 before they are translocated into the host cell (Khang et al. 2010; Giraldo et al. 2013). In addition to effectors, fungal antioxidation systems are essential to the success of biotrophic invasion by scavenging host reactive oxygen species (ROS) that otherwise trigger plant innate immunity (Huang et al. 2011; Marroquin-Guzman et al. 2017; Li et al. 2020; Rocha and Wilson, 2020). Despite being essential for disease progression, and although well described at the cellular level (Kankanala et al. 2007; Giraldo et al. 2013), very little is known at the molecular level about plant-fungal interfacial biology, including its integration with biotrophic growth (Sun et al. 2018). Consequently, important molecular details regarding the basic principles of plant-microbe interactions are obscured.

We seek to understand the metabolic strategies and molecular decision-making processes employed by *M. oryzae* to ensure metabolic homeostasis during biotrophy, when resources must be allocated between IH growth, maintaining interfacial integrity, and plant defense suppression. How fungal metabolic homeostasis is achieved during biotrophy is particularly intriguing when considering that genetic studies indicate fungal acquisition of host purines and amino acids is severely limited or non-existent during biotrophy (Wilson et al. 2012; Fernandez et al. 2013; Marroquin-Guzman et al. 2018). Recently, we uncovered a role for the *M. oryzae* target of rapamycin (TOR) signaling pathway in mediating biotrophic growth, maintaining biotrophic interface integrity (despite the plant origins of BIC and EIHM membranes) and ensuring correct effector deployment (Sun et al. 2018). Tor kinase is a conserved signaling hub that promotes growth in the presence of nutrients and energy and induces autophagy under starvation conditions (Loewith and Hall, 2011). Previously, in a forward genetics screen for TOR signaling components, we identified a vacuole membrane protein, Imp1, as a downstream target of TOR (Sun et al. 2018). Imp1 is not required for basal autophagy, but it is required for autophagy induction (Sun et al. 2018). *Δimp1* mutant strains penetrated leaf cuticles using functional appressoria, and developed IH with one or two branches, but IH failed to fill the first infected rice cell. Using the fluorescently labelled effector probes Pwl2-mCherry^NLS^ and Bas4-GFP (Khang et al. 2010) showed that *Δimp1* BICs and the EIHM eroded during early colonization (Sun et al. 2018). *Δimp1* biotrophic growth and interfacial integrity was restored by treatment with the TOR-independent autophagy inducer amiodarone hydrochloride (AM), while treatment of WT-infected rice cells with the autophagy inhibitor 3-methyladenine (3-MA) recapitulated the *Δimp1* phenotype. Thus, TOR-Imp1-autophagy signaling maintains membrane homeostasis via autophagosome formation and endomembrane trafficking to prevent interfacial erosion and support biotrophic growth (Sun et al. 2018).

To understand more about the molecular underpinnings of biotrophic growth, here, we screened genes for roles in maintaining biotrophic interfacial integrity by expressing Pwl2-mCherry^NLS^ and Bas4-GFP as fluorescent biotrophic membrane integrity probes in two mutant strains impaired for biotrophic growth: a mutant strain disrupted for the *M. oryzae* homologue of *RIM15* encoding a serine-threonine protein kinase acting on autophagy in parallel with TOR in yeast (Swinnen et al. 2006; Yorimitsu et al. 2007; Yang et al. 2010; Kim et al. 2021); and *Δasn1*, a previously characterized asparagine auxotrophic mutant (Marroquin-Guzman et al. 2018). Characterizing these mutant strains using a combination of live-cell imaging, leaf sheath infusions of glutamine-related metabolites and autophagy modifiers, genome-wide analyses and plate tests, revealed that biotrophic growth and interfacial membrane integrity requires cycles of autophagy-dependent TOR reactivation mediated by α-ketoglutarate, a previously unknown amino acid sufficiency signal (**Fig. 1a**). Thus, by delivering new insights into the fundamental nature and regulation of fungal biotrophic growth, and how fungal metabolism is integrated with host plant cell innate immunity suppression, this work lays a more solid foundation for comprehensively understanding pathogen colonization of living host cells at the systems biology level.

**Figure 1.**
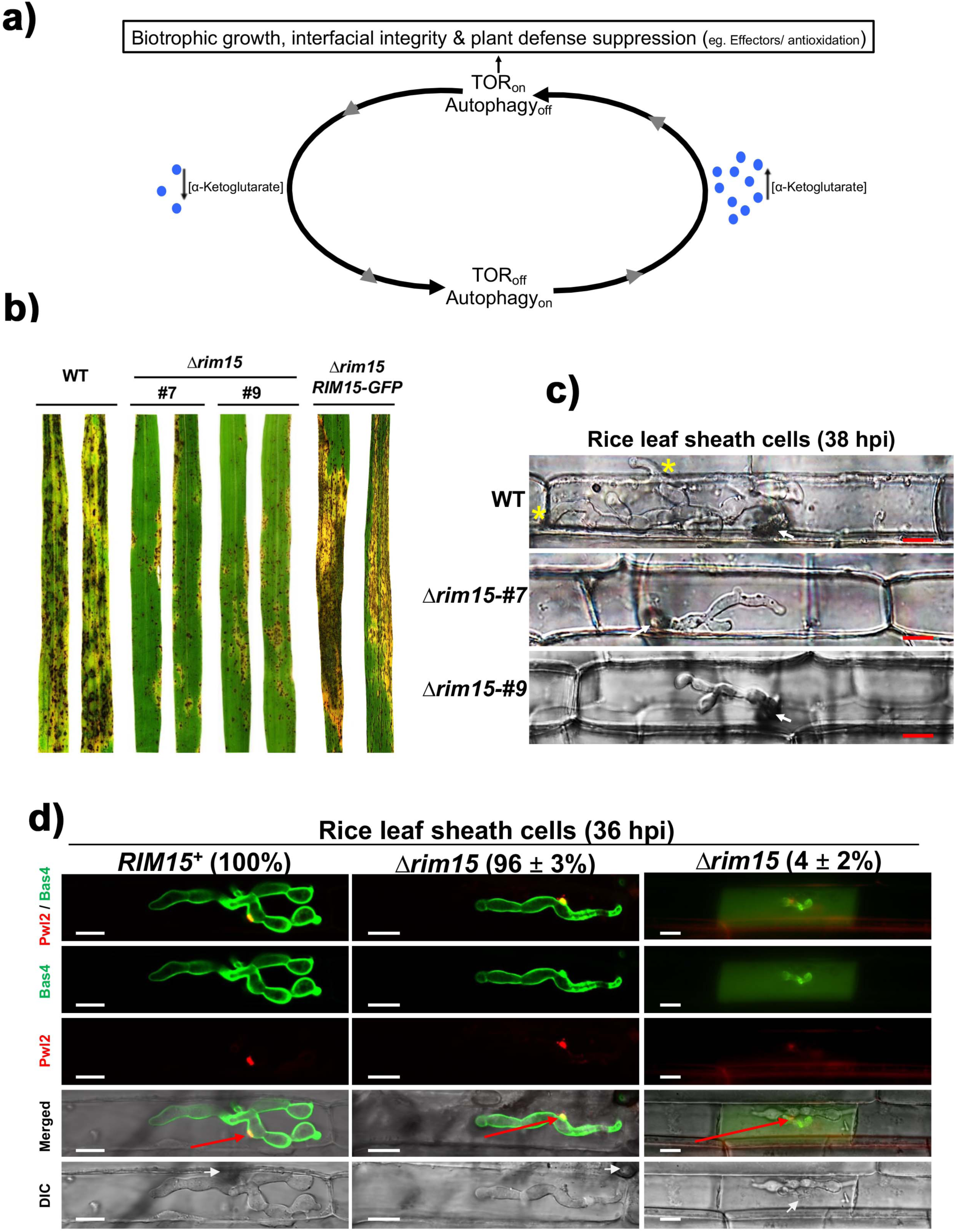
*M. oryzae RIM15* is required for biotrophic growth in host rice cells. **a**. Schematic overview of the major findings of this work. **b.** Rice blast disease symptoms of leaves infected with the indicated strains. Spores applied to 3-week-old rice seedlings of the susceptible cultivar CO-39 at a rate of 1 ×10^5^ spores ml^−1^. Images were taken at 120 hr post inoculation (hpi). **c.** Live-cell imaging at 38 hpi of detached rice leaf sheaths infected with the indicated strains showing how *Δrim15* strains are impaired for biotrophic growth. White arrows indicate appressorial penetration sites. Asterisks indicate movement of IH into neighbouring cells. Scale bar is 10 µm. **d.** Live-cell imaging at 36 hpi of detached rice leaf sheaths infected with strains expressing Pwl2-mCerry^NLS^ and Bas4-GFP shows how biotrophic membranes were intact in most rice cells (96 %) infected with the *Δrim15* mutant strain, but for a small subset (4 %), interfacial erosion is visible. White arrows indicate appressorial penetration sites. Red arrows indicate BICs in the merged channel for ease of viewing. Representative images and values are derived from observing 50 infected rice cells per strain, repeated in triplicate. Values are averages with standard deviation. DIC is differential interference contrast. Scale bar is 10 µm.

## Results

### The serine/threonine protein kinase Rim15 is required for biotrophic growth and interfacial membrane integrity

The serine/threonine protein kinase Rim15 is well characterized in yeast and functions to integrate signaling from the nutrient-sensing kinases TORC1, Sch9, PKA and Pho85-Pho80. Yeast Rim15 is required for starvation-induced autophagy in response to PKA and Sch9 inactivation, but it is not required for TORC1-dependent rapamycin-induced autophagy, suggesting PKA and Sch9 in yeast regulate autophagy in a pathway parallel to the TORC1 pathway (Swinnen et al. 2006; Yorimitsu et al. 2007; Yang et al. 2010; Kim et al. 2021). The blast fungus gene MGG_00345 (Dean et al. 2005) is the *M. oryzae RIM15* homologue. Using this sequence, we disrupted the *RIM15* coding region in our WT Guy11 isolate using homologous gene recombination to replace the first 1 kb of the *RIM15* coding sequence with the *ILV2* gene conferring sulphonylurea resistance (Sun et al. 2018). More than 10 *Δrim15-*carrying mutant strains were identified by PCR and two deletants, *Δrim15* #7 and #9, were initially characterized and found to be indistinguishable (**Fig 1b, c** and **Fig S1a-d**). *Δrim15* #7 was used as the recipient for the *RIM15-GFP* gene to generate the *Δrim15 RIM15-GFP* complementation strain (**Fig 1b**). *Δrim15* #7 was also the recipient of the pBV591 vector (Khang et al. 2010) to generate *Δrim15* expressing Pwl2-mCherry^NLS^ and Bas4-GFP (**Fig 1d**). Compared to the WT parental strain, *Δrim15* #7 and #9 deletion strains were both marginally reduced in radial diameter on complete media (CM), and both were reduced for sporulation on CM (**Fig S1a,b**). Sporulation was improved on oatmeal agar, enabling enough *Δrim15* spores to be harvested for downstream applications. Applying equal numbers of spores to 3-week-old seedlings of the susceptible CO-39 rice cultivar showed that compared to WT and the complementation strain, *Δrim15* #7 and #9 mutant strains were non-pathogenic, with both deletants producing only small, Type I lesions (Valent et al.1991) that did not produce spores. *Δrim15* #7 and #9 formed normal-looking appressoria that were melanized and inflated on artificial hydrophobic surfaces by 24 hpi (**Fig S1c**). *Δrim15* #7 and #9 appressoria formed at rates that were indistinguishable from WT on both hydrophobic surfaces and detached rice leaf sheath surfaces (**Fig S1d**), and *Δrim15* #7 and #9 appressoria penetrated rice leaf sheaths at rates comparable to WT (**Fig S1d**). However, although able to elaborate IH in the first infected cell, subsequent growth was curtailed compared to WT (**Fig 1c**). By 44 hpi, *Δrim15* #7 and #9 IH movement into adjacent cells (which occurs via plasmodesmata (Kankanala et al. 2007; Sakulkoo et al. 2018)) was rarely observed (**Fig S1d**) and if so, it only involved one or two IH strands that failed to grow far into the second infected cell (eg see **Fig 2b**). We conclude that *RIM15* is required for establishing extensive biotrophic growth in the first infected rice cell.

**Figure 2.**
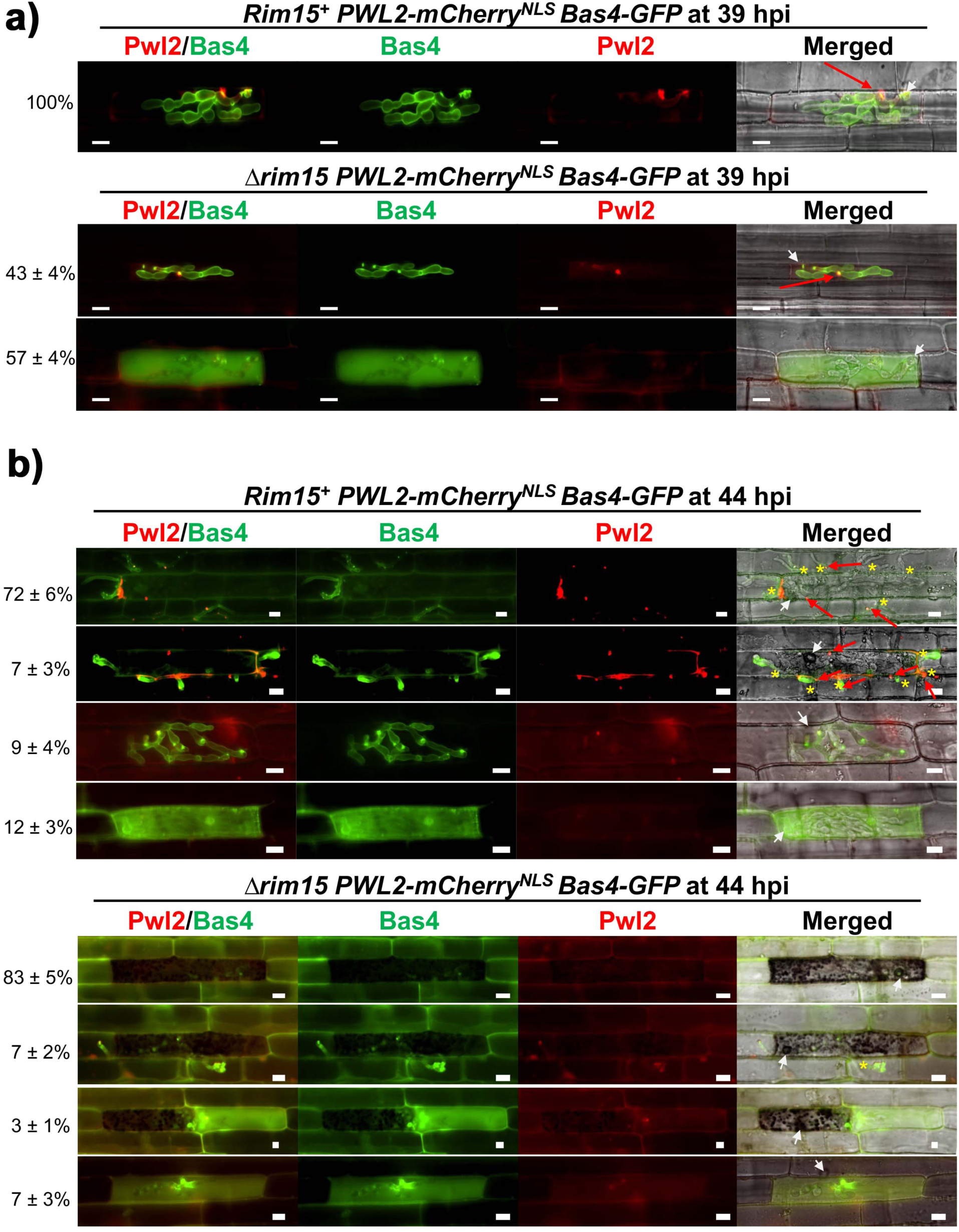
Loss of *RIM15* results in biotrophic interface erosion and the induction of strong plant defense responses. Live-cell imaging at 39 hpi (**a**) and 44 hpi (**b**) of detached rice leaf sheaths infected with strains expressing Pwl2-mCerry^NLS^ and Bas4-GFP shows how *Δrim15* biotrophic membranes erode over time. This membrane loss is accompanied by the induction of severe host plant cell responses that are not observed for *RIM15^+^*-infected rice cells at these time points. White arrows indicate appressorial penetration sites. Red arrows denote BICs. Asterisks indicate movement of IH into neighbouring cells. Representative images and values are derived from observing 50 infected rice cells per strain, repeated in triplicate. Values are averages with standard deviation. Scale bar is 10 µm.

We next employed our *Δrim15* strain carrying *PWL2-mCherry^NLS^* and *BAS4-GFP* to determine if attenuated *Δrim15* biotrophic growth impacted membrane integrity. By 36 hpi, in our *RIM15*^+^ strain expressing Pwl2-mCherry^NLS^ and Bas4-GFP, Pwl2 accumulated in the BIC and Bas4 accumulated in the apoplastic space to outline IH, as previously described (Khang et al. 2010; Giraldo et al. 2013; Marroquin-Guzman et al. 2017; Sun et al. 2018) (**Fig 1d**). For our *Δrim15* strain expressing Pwl2-mCherry^NLS^ and Bas4-GFP, although *Δrim15* IH was already impaired for growth by 36 hpi compared to WT, we nonetheless observed in most *Δrim15*-infected rice cells that, similar to WT, Pwl2 accumulated in prominent BICs and the EIHM was intact resulting in Bas4 accumulation in the apoplast (**Fig 1d**). However, about 4 % of *Δrim15*-infected rice cells displayed some Bas4 leakage into the host cytoplasm. By 39 hpi, when all WT-infected rice cells displayed intact BICs and EIHM, over half of all *Δrim15*-infected rice cells (n=50, repeated in triplicate) had lost BICs and were leaking Bas4 into the cytoplasm, indicating BIC and EIHM integrity was lost (**Fig 2a**). This is similar to what was observed for *Δimp1* strains (Sun et al. 2018). By 44 hpi (**Fig 2b**), all *Δrim15*-infected rice cells had lost their BICs and Bas4 was observed in the rice cytoplasm in some cases, but this was often obscured by very robust plant defense responses that manifested as black occlusions in the host cell. Such a strong host cell response was not observed for *Δimp1*-infected rice cells (Sun et al. 2018). Together, these results indicate that *RIM15* is not required for establishing IH growth and biotrophic interfacial membrane integrity but is, like *IMP1*, required for maintaining biotrophic interfacial membrane integrity and supporting sustained IH growth. Moreover, because, by 44 hpi, *Δrim15*-infected rice cells elicited an extremely robust host plant defense response (**Fig 2b**) – proceeded by the accumulation of host ROS at 30 hpi (**Fig S1e**) – that was not observed for *Δimp1*-infected rice cells (Sun et al. 2018), we concluded from these first experiments that Rim15 acted in the TOR-Imp1-autophagy signaling cascade, upstream of Imp1 but downstream of TOR, at a branch with a separate Rim15-controlled pathway required for mediating ROS scavenging and host innate immunity suppression.

In support of our hypothesis of an epistatic TOR-Rim15-Imp1-autophagy-biotrophic growth pathway (with a separate Rim15-host innate immunity suppression branch), **Fig 3a** shows that, like for *Δimp1* (Sun et al. 2018), the *Δrim15* mutant strain was impaired in autophagy and grew poorly on starvation media compared to WT. Also, like *Δimp1* (Sun et al. 2018), *Δrim15* biotrophic growth and interfacial membrane integrity was remediated at 44 hpi following application at 36 hpi of the autophagy stimulator AM to *Δrim15*-infected rice leaf sheaths (**Fig 3b**). However, unlike *Δimp1*, which was unresponsive to rapamycin treatment (Sun et al. 2018), *Δrim15* biotrophic interfacial integrity was remediated by rapamycin treatment (although rapamycin treatment attenuated biotrophic growth for both *RIM15^+^* and *Δrim15* strains) (**Fig 3b**). Therefore, we rejected our hypothesis of a linear relationship between TOR, Rim15 and Imp1 and instead proposed that, like in yeast (where Rim15 is also not required for rapamycin-induced responses), Rim15 acts on autophagy in parallel with the TOR-autophagy branch signaling pathway (**Fig 3c**).

**Figure 3.**
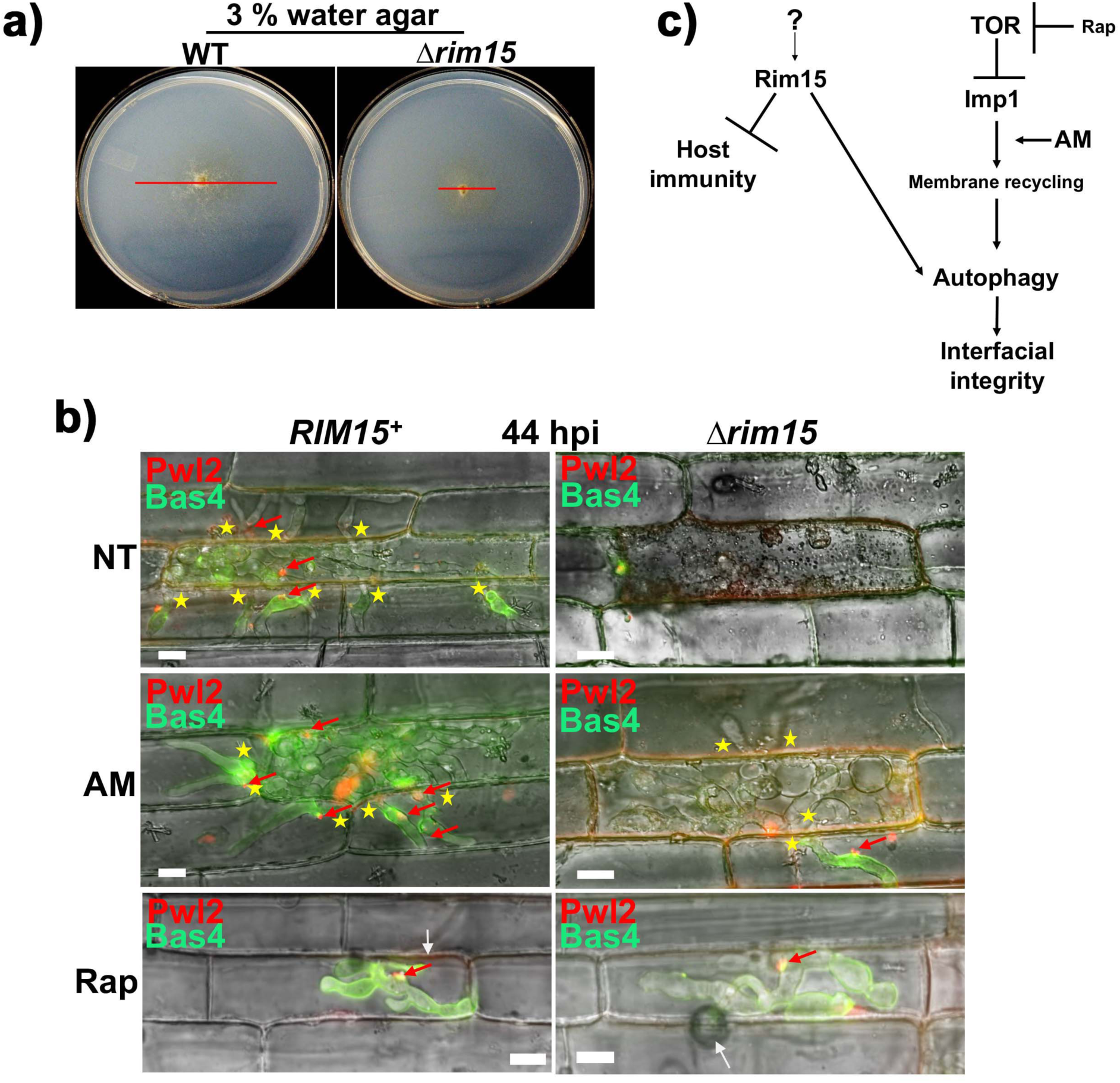
*RIM15* acts on autophagy in parallel with the TOR-Imp1-autophagy pathway. **a.** Plate tests showing impaired growth of the *Δrim15* mutant strain on 3 % water agar media without nutrients, compared to WT. Red bar indicates colony diameters. Images were taken at 12 dpi. **b.** Live-cell imaging of detached rice leaf sheaths infected with strains expressing Pwl2-mCherry^NLS^ and Bas4-GFP shows how amiodarone hydrochloride (AM) and rapamycin (Rap) treatment both remediated *Δrim15* biotrophic interfacial membrane integrity. AM but not rap treatment remediated *Δrim15* IH growth to adjacent cells. Leaf sheaths were treated with 1.5 μM AM or 10 μM Rap dissolved in 1% DMSO at 36 hpi and viewed at 44 hpi. White arrows indicate appressorial penetration sites. Red arrows denote BICs. Asterisks indicate movement of IH into neighbouring cells. All images are representative of 50 infected rice cells per strain, repeated in triplicate. Scale bar is 10 µm. NT is no treatment. Merged channel is shown. **c.** Model showing relationship between Rim15, TOR-Imp1 signaling and autophagy.

### Rim15 mediates autophagic flux during vegetative and biotrophic growth

To further elaborate the role of *RIM15* in biotrophic growth, we next focused on its connection to autophagy. To understand specifically how autophagy was affected in *Δrim15* strains, we generated a plasmid encoding GFP fused to the N-terminus of *M. oryzae* autophagy-related protein 8 (Atg8) and transformed it into WT and *Δrim15* #7 in order to perform the GFP-ATG8 processing assay for bulk autophagic activity (Klionsky et al. 2020) (**Fig 4a**). In this assay, autophagy activity results in autophagic bodies carrying GFP-Atg8 fusing with the vacuole, where Atg8 is degraded following lysis of the autophagic body, but GFP is more resistant to proteolysis and accumulates (Yorimitsu et al 2007; Klionsky et al. 2020; Kim et al. 2021). **Fig 4b** shows that, as expected, GFP accumulated in *RIM15^+^* vacuoles when vegetative mycelia were switched from nutrient-rich liquid CM to water, and no GFP-Atg8 accumulated in hyphal cytoplasm in water. In contrast, under the same growth conditions, *Δrim15* GFP- Atg8 accumulated little or no GFP in vacuoles in water, and GFP-Atg8 accumulated in mycelial cytoplasm in both CM and water. These results indicate that *Δrim15* strains are blocked in autophagy and that, like in yeast (Yorimitsu et al. 2007), Rim15 is required for autophagic flux but not for autophagy induction in response to nutrient starvation. During early biotrophic growth, at 28 hpi and 36 hpi, GFP accumulated in vacuoles in *RIM15^+^* GFP-Atg8 IH, indicating autophagy activity was high in WT at these very early infection time points (**Fig 4c,d**). This high autophagy activity in WT had not been shown before in *M. oryzae* IH and was unexpected at such early time points. In contrast, at 28 hpi and 36 hpi, GFP-Atg8 localized to the cytoplasm in *Δrim15* and did not accumulate in vacuoles (**Fig 4c,d**). Therefore, WT is undergoing autophagy during early biotrophic infection, but autophagic activity is blocked in *Δrim15*. Note that 36 hpi corresponds in *Δrim15* to mostly intact biotrophic interfaces (**Fig 1d**) and is on the cusp of the catastrophic interfacial membrane integrity loss seen at 39 hpi and later for this mutant strain (**Fig 2**).

**Figure 4.**
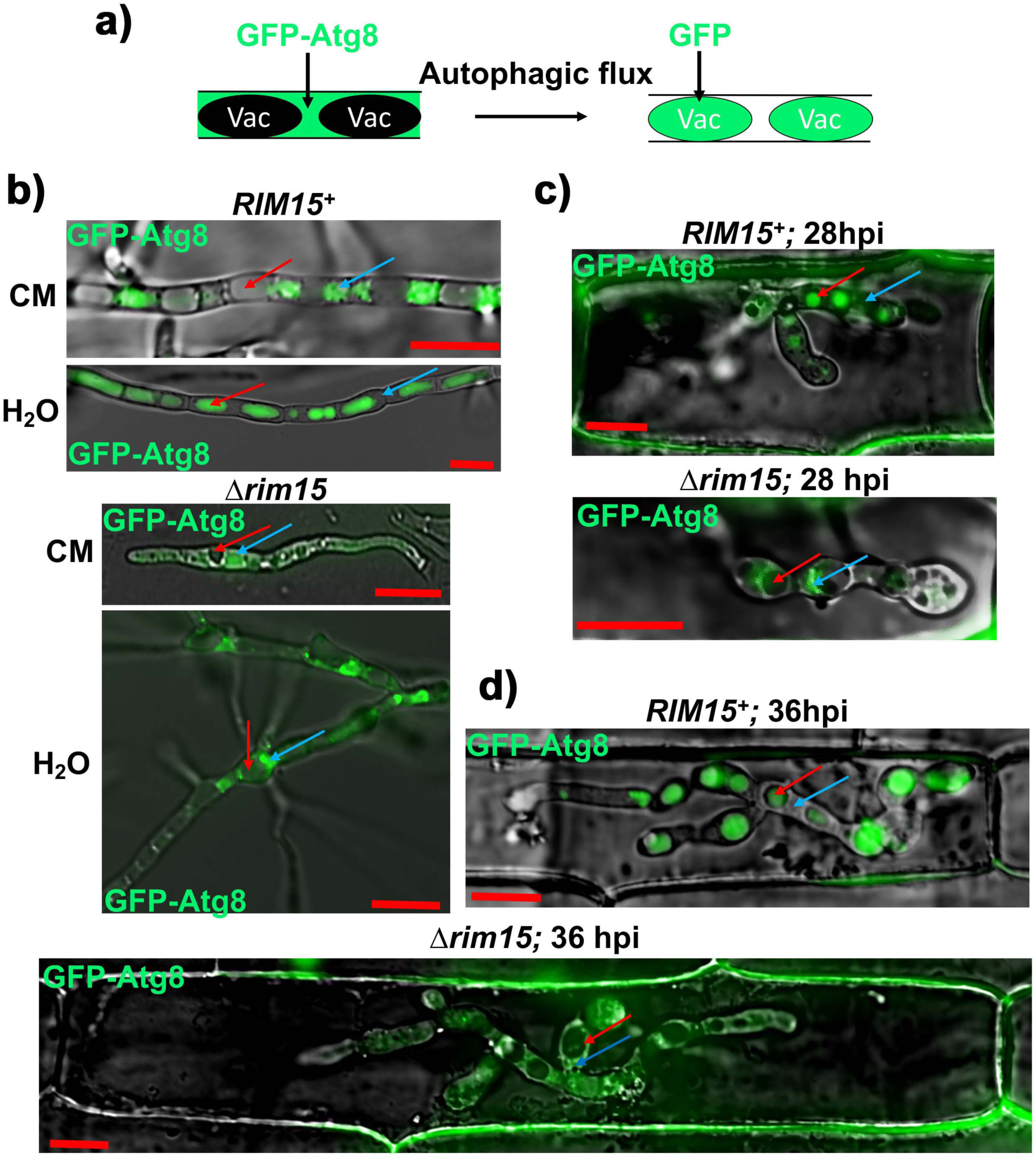
Rim15 is required for autophagic flux in vegetative hyphae and IH. **a.** Schematic of the GFP-Atg8 bulk autophagy activity assay. Vac are vacuoles. **b-d.** Micrographs showing how autophagy activity is attenuated in *Δrim15* strains compared to WT. Examples of vacuoles are indicated with red arrows, examples of cytoplasm are indicated with blue arrows. Bar is 10 µm. Merged channel is shown. **b.** Mycelia of the indicated strains expressing GFP-Atg8 were inoculated in liquid CM medium for 42 hr before harvesting. Mycelia was washed with water and inoculated into fresh liquid CM or water, as indicated, for 3.5 hr before imaging. **c, d.** Live-cell imaging of detached rice leaf sheaths infected with strains expressing GFP-Atg8 at the indicated time points. All images are representative of 50 infected rice cells per strain, repeated in triplicate, and obtained using a Nikon Eclipse Ni-E upright microscope at 28 hpi (**c**) and 36 hpi (**d**).

### Rim15 is required for autophagic cycling *in planta*

Live-cell imaging at time points later than 36 hpi showed that GFP accumulated in *Δrim15* vacuoles at 40 hpi and 44 hpi, suggesting that autophagic flux was delayed but not abolished in the mutant strain (**Fig 5a**). GFP accumulated in *RIM15^+^* vacuoles at 44 hpi (**Fig 5a**), indicating that autophagy is active as the fungus is moving to neighbouring cells, which is consistent with our earlier findings that AM treatment promotes cell-to-cell invasion (Sun et al. 2018). However, intriguingly, we observed that autophagic flux was reduced in the *RIM15^+^* GFP-Atg8 strain at 40 hpi, with most of the GFP signal accumulating as GFP- Atg8 in the cytoplasm (**Fig 5a**). To understand autophagy dynamics better, we quantified the percentage of *M. oryzae* vacuoles accumulating GFP, counted in fifty infected rice cells per time point (repeated in triplicate). This revealed that in WT, autophagy activity levels cycle during biotrophic growth, but autophagic cycling was abolished in *Δrim15* (**Fig 5b**). We conclude that during biotrophy, Rim15 is required for autophagic cycling, a hitherto unknown phenomenon in *M. oryzae* IH.

**Figure 5.**
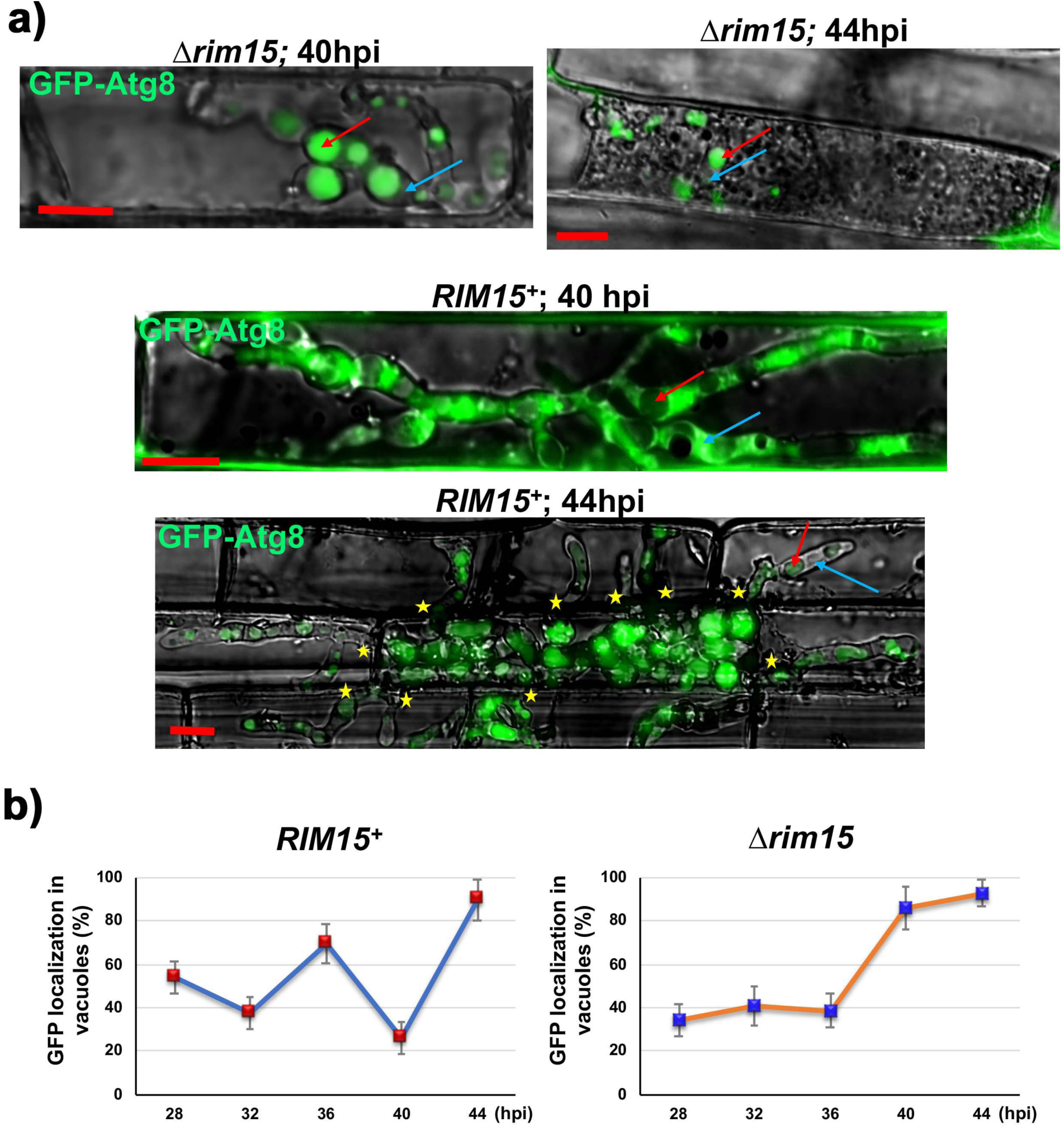
Autophagic cycling in IH is Rim15-dependent. **a.** Live-cell imaging of detached rice leaf sheaths infected with strains expressing GFP-Atg8 shows how in *Δrim15* strains at 40 hpi and 44 hpi, autophagic flux is remediated relative to earlier time points. In contrast, in *RIM15^+^* strains, autophagic activity is reduced at 40 hpi relative to earlier time points but increases again at 44 hpi. All images are representative of 50 infected rice cells per strain, repeated in triplicate. Examples of vacuoles are indicated with red arrows, examples of cytoplasm are indicated with blue arrows. Asterisks indicate movement of IH into neighbouring cells. Bar is 10 µm. Merged channel is shown. **b.** Quantification of GFP localization in vacuoles during IH growth shows how autophagy activity cycles in WT, but autophagic cycling is abolished in *Δrim15*. Values are the average number of IH vacuoles accumulating GFP in 50 infected rice cells as a percentage of the total number of vacuoles, repeated in triplicate. Error bars are standard deviation.

### Autophagic cycling is physiologically relevant to biotrophic colonization

Previously, we had shown that treating WT-infected rice cells with the autophagy inhibitor 3-MA at 36 hpi had, by 44 hpi, aborted IH growth and abolished cell-to-cell movement, thereby phenocopying the loss of *IMP1* (Sun et al. 2018). To determine if the autophagic cycling observed in **Fig 5b** had physiological relevance during infection, we treated WT-infected rice cells with 3-MA at 36, 40 and 44 hpi corresponding to two peaks and a trough of autophagy activity in WT (**Fig 5b**). When viewed at later time points, we found that in all cases, biotrophic growth was attenuated following 3-MA treatment relative to the untreated WT control (**Fig S2**). In many cases, a plant response was also visible (**Fig S2**). Therefore, blocking autophagy during active autophagic cycling, including after IH has penetrated to the second infected rice cell, aborted further IH growth, suggesting cycles of autophagy activity are required for IH growth throughout biotrophy.

### *M. oryzae* Rim15 does not accumulate in the nucleus

In yeast, under nutrient starvation and other stress conditions, Rim15 translocates to the host nucleus in a manner dependent on its cytoplasmic retention by Sch9 (Pedruzzi et al. 2003; Wanke et al. 2008; Kim et al. 2021). However, *M. oryzae* Rim15 appears constitutively localized to the cytoplasm and does not accumulate in the nucleus. Although Rim15 carries predicted nuclear localization signals (according to PSORTII), and even though our *Δrim15 RIM15-GFP* complementation strain expressed a functional Rim15-GFP protein as evidenced by both its ability to restore pathogenicity in the *Δrim15* background (**Fig 1b**) and its remediation of *Δrim15* colony growth on plates (**Fig S3a**), *M. oryzae* Rim15-GFP did not accumulate in the nucleus under any tested growth condition, nor did the amount of Rim15-GFP in the cytoplasm diminish. In *Δrim15 RIM15-GFP* vegetative hyphae, Rim15-GFP was cytoplasmic under both nutrient-rich and nutrient-starvation shake conditions (**Fig S3b**). Rim15-GFP was cytoplasmic in *Δrim15 RIM15-GFP* IH from 28 hpi until at least 44 hpi (**Fig S3c**), time points that correspond with autophagic cycling (**Fig 5b**) and cell-to-cell movement of IH. Expressing *RIM15-GFP* in a strain of *M. oryzae* carrying histone H1 fused to RFP (Saunders et al. 2010) showed that Rim15-GFP did not co-localize with nuclei during biotrophy (shown for 36 hpi in **Fig S3d**). Therefore, *M. oryzae* Rim15-GFP functions similarly to yeast Rim15 but does not accumulate in the nucleus in order to mediate autophagy or biotrophic growth, at least under our tested conditions and time points. Differences in ScRim15 and MoRim15 localization may reflect evolutionary changes in response to the different lifestyles of these two widely separated groups of fungi.

### Rim15 is required for glutaminolysis

To better understand the function of Rim15 in biotrophy, we next turned our attention to RNAseq data obtained at 36 hpi (ie before *Δrim15* biotrophic interfacial membrane loss) from *Δrim15*-, WT- and mock-infected rice leaf sheaths (**Table S1**). Our attention was drawn to genes encoding enzymes of the GS-GOGAT cycle (Mora, 1990) that were altered in expression in *Δrim15* IH at 36 hpi compared to WT (**Fig 6a**). The GS-GOGAT cycle is required for ammonium assimilation and the interconversion of glutamine, glutamate and *α*-ketoglutarate, and also for glutaminolysis, the use of glutamine or glutamate as carbon sources through conversion to *α*-ketoglutarate (**Fig 6b**). We focused on this pathway because we had plate test data in hand showing that *Δrim15* strains were attenuated for glutaminolysis (**Fig 6b**). Specifically, *Δrim15* growth was severely restricted compared to WT on minimal media containing glutamine or glutamate as the sole carbon and nitrogen source, but *Δrim15* grew better on minimal media containing glutamine or glutamate as sole nitrogen sources with glucose as the preferred carbon source.

**Figure 6.**
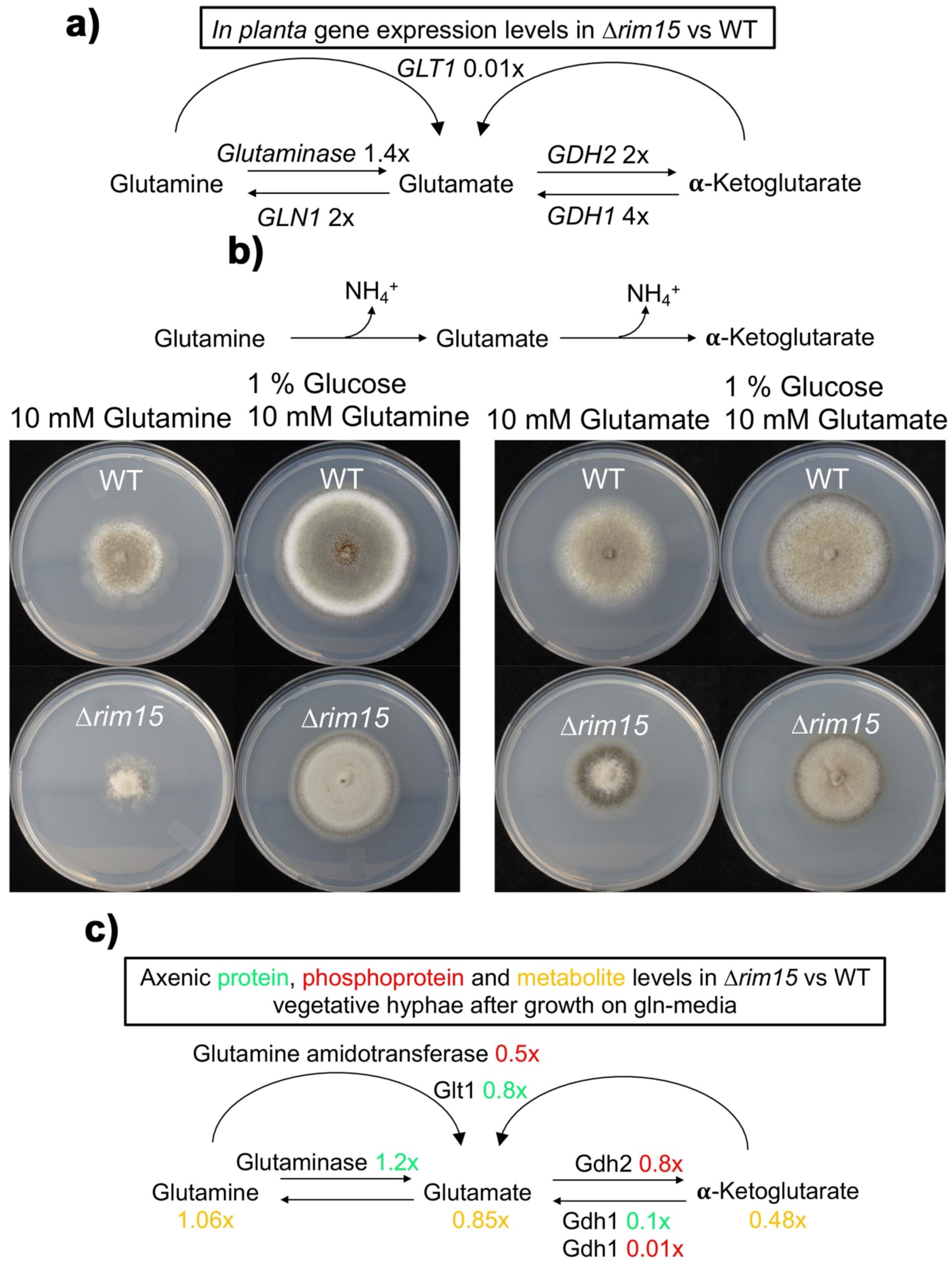
*Δrim15* is perturbed in glutaminolysis and *α*-ketoglutarate homeostasis compared to WT. **a.** Simplified schematic of the GS-GOGAT cycle summarizing RNAseq data (**Table S1**) from 15 infected detached rice leaf sheaths per strain, repeated in triplicate and compared to the mock infection control, showing how GS-GOGAT cycle gene expression changes in *Δrim15* IH at 36 hpi are misregulated compared to WT. *GLN1* (MGG_14279) encodes glutamine synthetase; *GLT1* (MGG_07187) encodes glutamate synthase (GOGAT); *GDH1* (MGG_08074) encodes NADP-specific glutamate dehydrogenase; *GDH2* (MGG_05247) encodes NAD-specific glutamate dehydrogenase (Mgd1) (Marroquin-Guzman and Wilson, 2015). Glutaminase is encoded at MGG_07512. GS-GOGAT cycle gene transcripts with equal abundances in both strains are not shown. Vaules are fold changes in *Δrim15*. **b.** Plate tests show how glutaminolysis (the use of glutamine or glutamate as a carbon source) is impaired in *Δrim15* compared to WT. **c.** Schematic summarizing GS-GOGAT cycle-related proteomic, phosphoproteomic and metabolomic data, averaged from three mycelial samples per strain grown in liquid minimal media with 0.5 mM glutamine as the sole carbon and nitrogen source (**Table S2-S5**), showing how *Δrim15* strains are perturbed for the GS-GOGAT cycle and glutaminolytic metabolite homeostasis. Strains were grown in CM for 42 hr before switching to 0.5 mM glutamine media for 16 hr. GS-GOGAT cycle enzymes with equal abundances in both strains are not shown. Values are fold changes in *Δrim15*.

The RNAseq and plate test data both pointed to GS-GOGAT cycle and glutaminolysis perturbations in *Δrim15*. To better understand the connection between Rim15 and glutaminolysis, we extracted proteins and metabolites for (phospho)proteomic and metabolomic analyses from WT and *Δrim15* mycelia grown in liquid minimal media with 0.5 mM glutamine as the sole carbon and nitrogen source. Proteomic (**Table S2**), quantitiative phosphoproteomic (**TableS3)** and metabolomic (**Table S3**) analyses on three independent mycelial sampes per strain showed that, to a greater or lesser extent, a number of proteins and metabolites associated with the GS-GOGAT pathway and glutaminolysis were perturbed in *Δrim15* compared to WT during growth under glutaminolytic conditions (summarized in **Fig 6c**). Most strikingly, *Δrim15* perturbations to the GS-GOGAT cycle resulted in a shift in glutamine/ glutamate homeostasis and a 2-fold decrease in *α*-ketoglutarate steady state levels in *Δrim15* vegetative hyphae compared to WT. Thus, glutaminolysis and the GS-GOGAT cycle are perturbed in *Δrim15*, affecting *α*-ketoglutarate homeostasis.

### a-ketoglutarate treatment remediated *Δrim15* biotrophic growth

Based on the previous results, we hypothesized that Rim15-mediated glutaminolysis and the GS-GOGAT cycle was required for biotrophy, and that the loss of *α*-ketoglutarate homoestasis in *Δrim15* was physiologically relevant to the infection process. We thus speculated that restoring *α*-ketoglutarate levels in *Δrim15* would improve infection. To test this, before attempting to manipulate *α*-ketoglutarate levels by pharmacological or genetic means, we first asked whether treating WT and *Δrim15*-infected rice cells with *α*-ketoglutarate would affect biotrophic growth. Unexpectedley, **Fig 7a** shows that treating *Δrim15*-infected rice cells at 36 hpi with 10 mM *α*-ketoglutarate (as the cell-permeable dimethyl *α*-ketoglutarate (DMKG) analog) completely remediated the *Δrim15* mutant phenotype by 44 hpi. Compared to the untreated *Δrim15* control, treatment with *α*-ketoglutarate suppressed the host defense response in *Δrim15*-infected rice cells and promoted cell-to-cell movement of *Δrim15* IH. *Δrim15* IH in adjacent host cells were outlined with Bas4, indicating an intact EIHM, and new BICs formed at IH tips. *Δrim15* IH development and spread in *α*-ketoglutarate-treated rice leaf sheaths was indistingushable at 44 hpi from *α*-ketoglutarate-treated *RIM15^+^* IH, which was itself indistinguishable from *RIM15^+^* IH growth and development in untreated rice cells (**Fig 7a**). Using GFP-Atg8 expressing strains, **Fig 7b** shows that compared to the untreated *Δrim15* control, *α*-ketoglutarate treatment at 36 hpi suppressed *Δrim15* autophagy by 44 hpi, leading to GFP-Atg8 accumulation in the cytoplasm and little GFP accumulation in vacuoles. Similarly, *α*-ketoglutarate treatment of *RIM15^+^* infected cells at 36 hpi suppressed autophagy at 44 hpi, resulting in GFP-ATG8 accumulation in the cytoplasm and no GFP in vacuoles even though *RIM15^+^* IH was extensively moving cell-to-cell by this time (**Fig S4**). This indicates that autophagy is not required for cell-to-cell spread in the presence of exogenous *α*-ketoglutarate. We conclude that *α*-ketoglutarate treatment remediates *Δrim15* biotrophic growth and suppresses autophagy in *RIM15^+^* and *Δrim15* IH.

**Figure 7.**
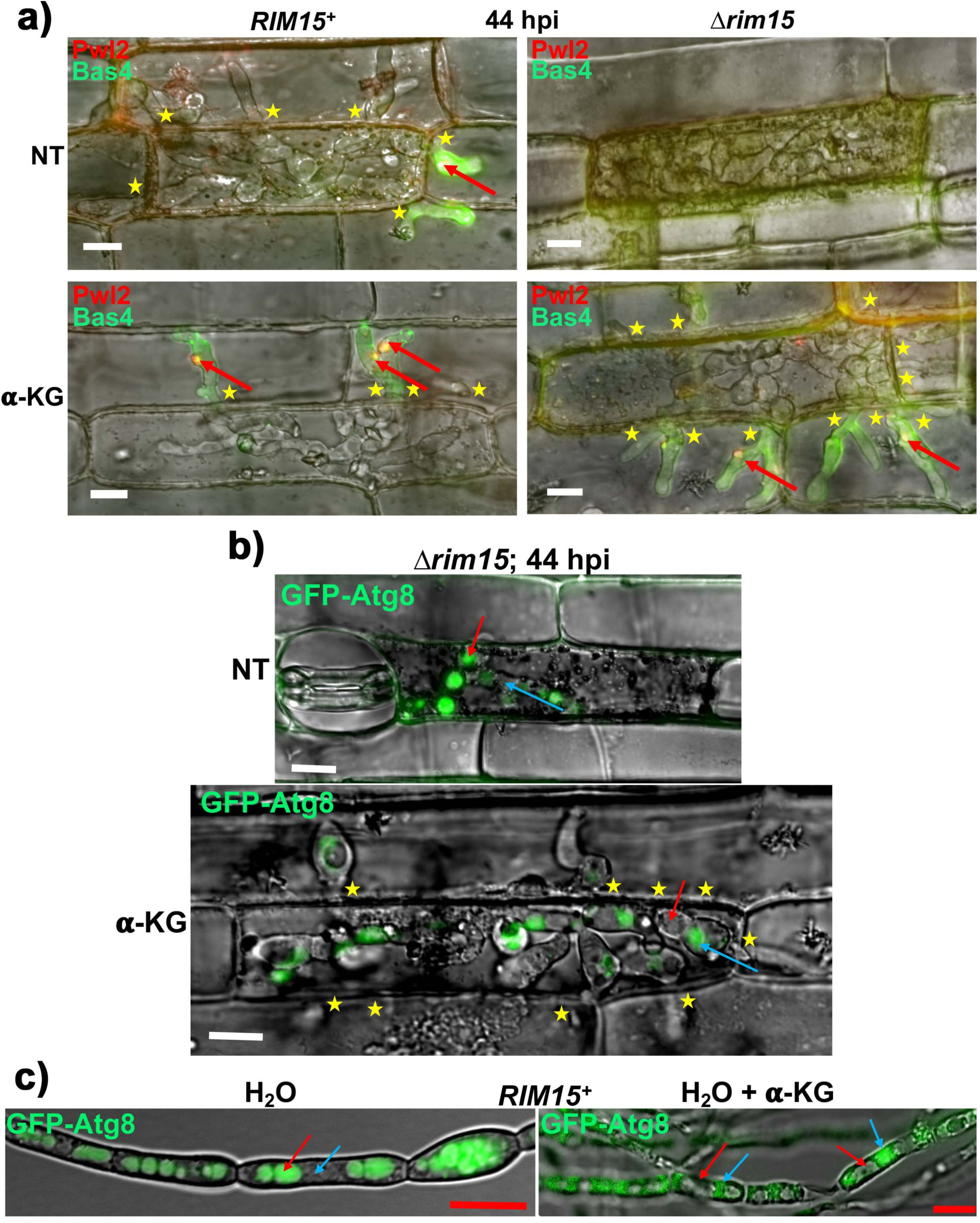
*α*-ketoglutarate treatment remediates *Δrim15* biotrophic growth defects and suppresses autophagy. **a.** Live-cell imaging of detached rice leaf sheaths infected with strains expressing Pwl2-mCherry^NLS^ and Bas4-GFP shows how treatment with 10 mM of the cell-permeable *α*-ketoglutarate (*α*- KG) analogue dimethyl *α*-ketoglutarate (DMKG) remediated *Δrim15* biotrophic growth and restored biotrophic interfacial membrane integrity. Red arrows indicate BICs. Asterisks indicate movement of IH into neighbouring cells. NT is no treatment. Bar is 10 µm. **b.** Micrographs show how 10 mM *α*-KG treatment (as DMKG) suppresses autophagy activity in *Δrim15* at 44 hpi. Examples of vacuoles are indicated with red arrows, examples of cytoplasm are indicated with blue arrows. Asterisks indicate movement of IH into neighbouring cells. NT is no treatment. Bar is 10 µm. **a, b.** 10 mM *α*-ketoglutarate (*α*- KG, as DMKG) was applied to detached rice leaf sheaths at 36 hpi, and images were taken at 44 hpi. All images are representative of 50 infected rice cells per strain, repeated in triplicate. Merged channel is shown. **c**. Micrograph showing that, although exposure to water alone for 4 hr induced autophagy in WT and GFP accumulated in vacuoles, in contrast, exposure to 10 mM *α*-KG (as the cell-permable DMKG analogue) in water for 4 hr was sufficient to suppress autophagy, leading to GFP-ATG8 accumulation in the cytoplasm. Mycelia were first grown in liquid CM for 42 hr before switching into the indicated treatments. Examples of vacuoles are indicated with red arrows, examples of cytoplasm are indicated with blue arrows. Bar is 10 µm. Merged channel is shown.

### *α*-ketoglutarate is an autophagy suppressing signal

*α*-ketoglutarate is only a carbon source and could not support fungal growth on its own. Thus, we hypothesized that *α*-ketoglutarate was a signal promoting IH growth and suppressing autophagy. Moreover, because these two cellular processes are indicative of TOR activity, we hypothesized that *α*-ketoglutarate acted by triggering TOR. Such a signaling role for *α*-ketoglutarate would account for both the remediation of *Δrim15* biotrophic growth following *α*-ketoglutarate treatment (as DMKG) (**Fig 7a**), and the autophagy suppression we observed *in planta* in *α*-ketoglutarate-treated WT IH (**Fig S4**). To test for this role, we switched *RIM15^+^* GFP-Atg8 vegetative mycelia from liquid CM shaking cultures (where GFP-Atg8 was mostly in cytoplasm) to water (where GFP accumulated in vacuoles) and into water + 10 mM *α*-ketoglutarate (as the cell-permable DMKG analog). Whereas switching vegetative hyphae into water alone triggered autophagy (resulting in no GFP-ATG8 in cytoplasm and GFP accumulation in vacuoles), the addition of *α*-ketoglutarate to water was sufficient to suppress autophagy (resulting in GFP-ATG8 accumulation in cytoplasm and no GFP accumulation in vacuoles), even though other nutrients including a nitrogen source were absent (**Fig 7c**). Thus, *α*-ketoglutarate is an autophagy suppressing signal. More specifically, because *α*-ketoglutarate is generated from the glutaminolysis of carbon- and nitrogen-containing amino acids (a process pertubed in *Δrim15*), we conclude that *α*-ketoglutarate is a carbon and nitrogen sufficiency signal that triggers TOR signaling, thereby suppressing autophagy and promoting biotrophic growth.

### Treatment with glutamine and glutamate, but not ammonium or asparagine, remediated *Δrim15* biotrophic growth in a dose-dependent manner

Because *Δrim15* was impaired in glutaminolysis and remediated by *α*-ketoglutarate, we tested whether other glutaminolysis-related metabolites remediated *Δrim15* biotrophic growth. Following treatment at 36 hpi, we found that by 44 hpi, cell-to-cell spread of *Δrim15* IH was remediated by 10 mM glutamine and 10 mM glutamate treatment, but not by 10 mM ammonium (NH_4_^+^) treatment (**Fig S5**). Treatment with 10 mM asparagine, which is not a glutaminolytic intermediate nor part of the GS-GOGAT cycle, also did not remediate *Δrim15* biotrophic growth (**Fig S5**). Furthermore, remediation by glutamine was dose-dependent, such that 1 mM or less of exogenously applied glutamine did not remediate *Δrim15* IH growth (**Fig S6**). Note that the cell-permeable *α*-ketoglutarate analog DMKG could not be reliably used to assess the effects of low concentrations of *α*-ketoglutarate on *Δrim15* IH biotrophic growth. Treatment with glutamine, glutamate or asparagine at 36 hpi had no effect on *RIM15^+^* IH growth by 44 hpi (**Fig S5**), but NH_4_^+^ treatment partialy inhibited *RIM15^+^* IH growth in the the second infected rice cells following IH spread, and NH_4_^+^ treatment also led to partial loss of *RIM15^+^* biotrophic membranes in the second infected cells. Together, our interpretation of these results is that exogenous treatments with high levels of exogenous glutamine or glutamate can remediate glutaminolytic homeostasis and drive *α*-ketoglutarate production, resulting in remediation of *Δrim15* IH biotrophic growth. Furthermore, our results indicate that the acquisition of these amino acids from the host plant cell by *Δrim15* must be limited. We can extend this principle to WT because 10 mM *α*-ketoglutarate treatment suppressed WT autophagy at 44 hpi (**Fig S4a**), suggesting WT access to *α*-ketoglutarate-generating host nutrients is likewise severely limited during biotrophy.

### The GS-GOGAT cycle is required upstream of *α*-ketoglutarate signaling for IH growth and biotrophic interfacial membrane integrity

Pharmacological evidence that the GS-GOGAT cycle is required for biotrophic growth and interfacial membrane integrity is shown in **Fig 8**. Treatment of WT-infected detached rice leaf sheaths at 36 hpi with the irreversible glutamine synthetase (GS) inhibitor methionine sulfoximine (MSO), or with the GOGAT inhibitor azaserine (AZS), had, in both cases by 44 hpi, resulted in attenuated growth and the loss of the EIHM, resulting in Bas4 in the rice cytoplasm. Unlike in *Δrim15*-infected rice cells, MSO- and AZS-treated WT-infected rice cells retained a small BIC, although in both cases Pwl2 had leaked into the rice cytoplasm, which was seen clearly in the red channel in **Fig 8**, indicating that BIC integrity was compromised compared to the untreated control. AZS-induced attenuation of IH growth and biotrophic interfacial membrane integrity was remediated by addition of the GOGAT product glutamate, while MSO toxicity was remediated by addition of the GS product glutamine, thus confirming AZS and MSO, repectively, as specific fungal GS-GOGAT cycle inhibitors *in planta*. Furthermore, *α*-ketoglutarate treatment could remediate AZS-induced biotrophic growth defects. Together, these results highlight the importance of the GS-GOGAT cycle to biotrophic growth and interfacial membrane integrity, likely as a means to supply *α*-ketoglutarate for downstream signaling purposes.

**Figure 8.**
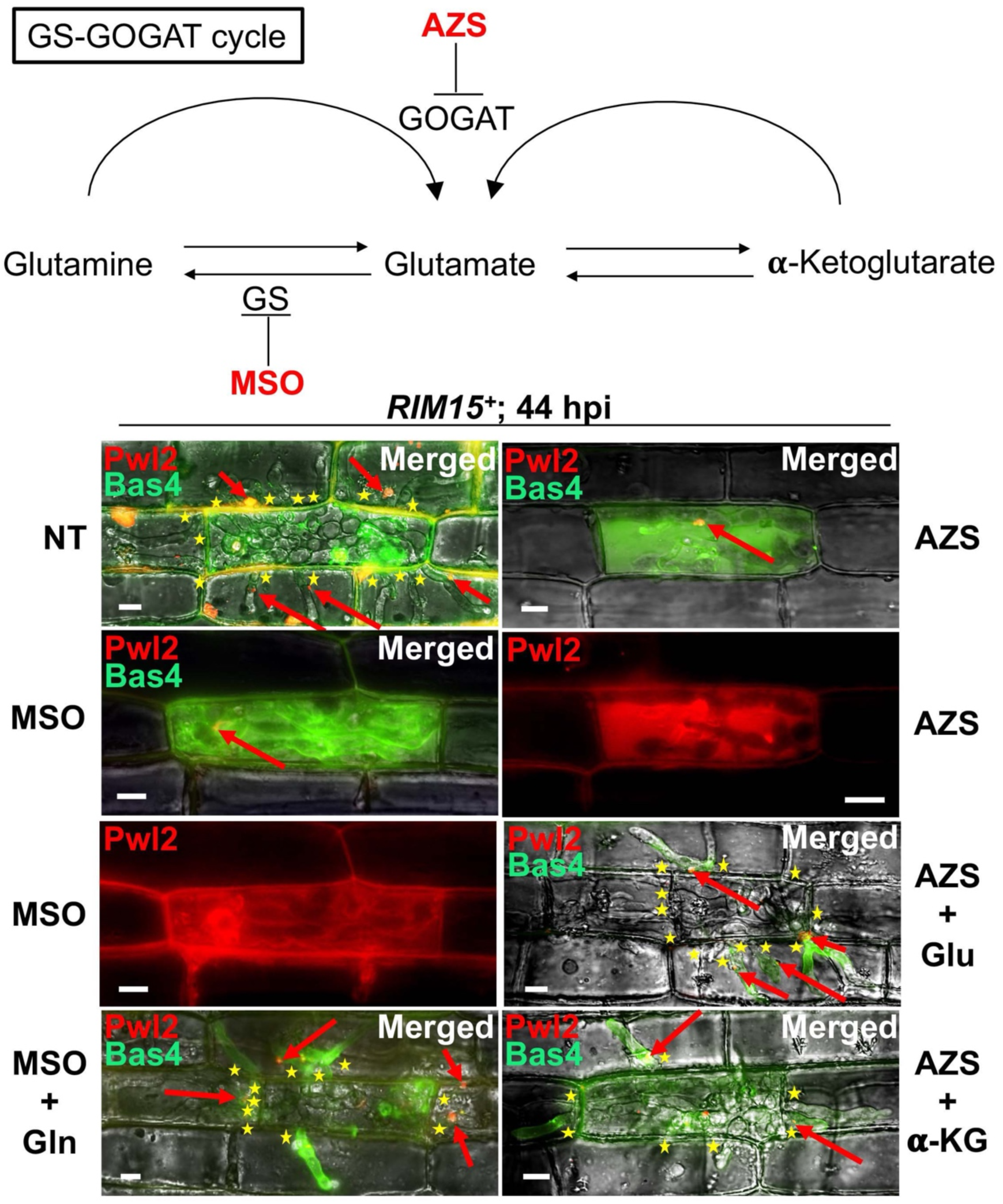
The GS-GOGAT cycle is required for biotrophic growth and biotrophic interfacial integrity. Live-cell imaging using WT expressing Pwl2-mCherry^NLS^ and Bas4-GFP shows how treating *RIM15^+^*-infected rice cells at 36 hpi with 1 mM L-methionine sulfoximine (MSO), the glutamine synthetase inhibitor, or with 2 mM azaserine (AZS), the GOGAT inhibitor, had, by 44 hpi, abolished biotrophic growth and thus phenocopied *Δrim15*. For MSO + Gln treatments, 1 mM MSO was first applied to the detached rice leaf sheath at 36 hpi and then replaced by 1 mM MSO + 10 mM glutamine at 40 hpi before imaging at 44 hpi. For AZs + Glu and AZS + *α*-KG, 2 mM AZS was first applied to the detached rice leaf sheath at 36 hpi then replaced at 40 hpi by 2 mM AZS + 10 mM glutamate or 2 mM AZS + 10 mM *α*-ketoglutarate (as the cell-permable DMKG analog) before imaging at 44 hpi. All images are representative of 50 infected rice cells per strain, repeated in triplicate. BICs are indicated with a red arrow. Asterisks indicate movement of IH into neighbouring cells. NT is no treatment. Bar is 10 µm.

### Autophagy-dependent biotrophic growth and interfacial membrane integrity requires amino acid prototrophy upstream of the *α*-ketoglutarate signal

We next found that IH growth and biotrophic interfacial membrane integrity was dependent on *M. oryzae* amino acid prototrophy. In a previous study, we showed that an *M. oryzae* asparagine auxotroph lacking asparagine synthetase, *Δasn1*, could make functional appressoria and elaborate IH but failed to establish extensive biotrophic growth in the first infected rice cell, indicating that asparagine and perhaps other amino acids were not accessible to this mutant strain during biotrophy (Marroquin-Guzman et al. 2018). Here, we asked, what happens to *Δasn1* biotrophic interfaces when IH growth is aborted? To address this question, we knocked out the *ASN1* gene in our strain carrying pBV591 to generate the mutant strain *Δasn1 PWL2-mCherry^NLS^ Bas4-GFP*. Live-cell imaging showed that by 44 hpi, this mutant strain was attenuated for biotrophic growth and had lost EIHM integrity, as evidenced by Bas4 in the rice cytoplasm (**Fig 9a**). The *Δasn1* BIC was disrupted at 44 hpi, although not entirely eradicted, but Pwl2 was observed in the rice cyctoplasm, suggesting BIC integrity was compromised and leaking Pwl2 (**Fig 9a**). *Δasn1* IH growth was partially restored (ie *Δasn1* IH did not fill the first infected cell), and cell-to-cell movement and biotrophic membrane integrity was fully restored, following treatment of infected rice leaf sheaths with 10 mM asparagine at 36 hpi (**Fig 9b**). Note that in an earlier report (Marroquin-Guzman et al. 2018), 10 mM asparagine added at 0 hpi to spore suspensions did not remediate biotrophic growth, indicating that the timing and means of application of remediating treatments for IH growth mutants is critical. Like for *Δrim15*, treatment with *α*-ketoglutarate remediated biotrophic interfacial membrane integrity and cell-to-cell movement, although this also occurred without filling the first infected rice cell, suggesting *α*-ketoglutarate could not completely overcome the loss of *de novo* asparagine biosynthesis with regards to IH growth (**Fig 9b**). However, unlike for *Δrim15* (and *Δimp1*), IH growth and biotrophic interfacial membrane integrity was not remediated by AM treatment (**Fig 9b**). Together, these results demonstrate that amino acid prototrophy is required for autophagy-mediated remediation of biotrophic growth and interfacial membrane integrity in IH growth-deficient mutants, but this requirement can be by-passed by sufficient *α*-ketoglutarate.

**Figure 9.**
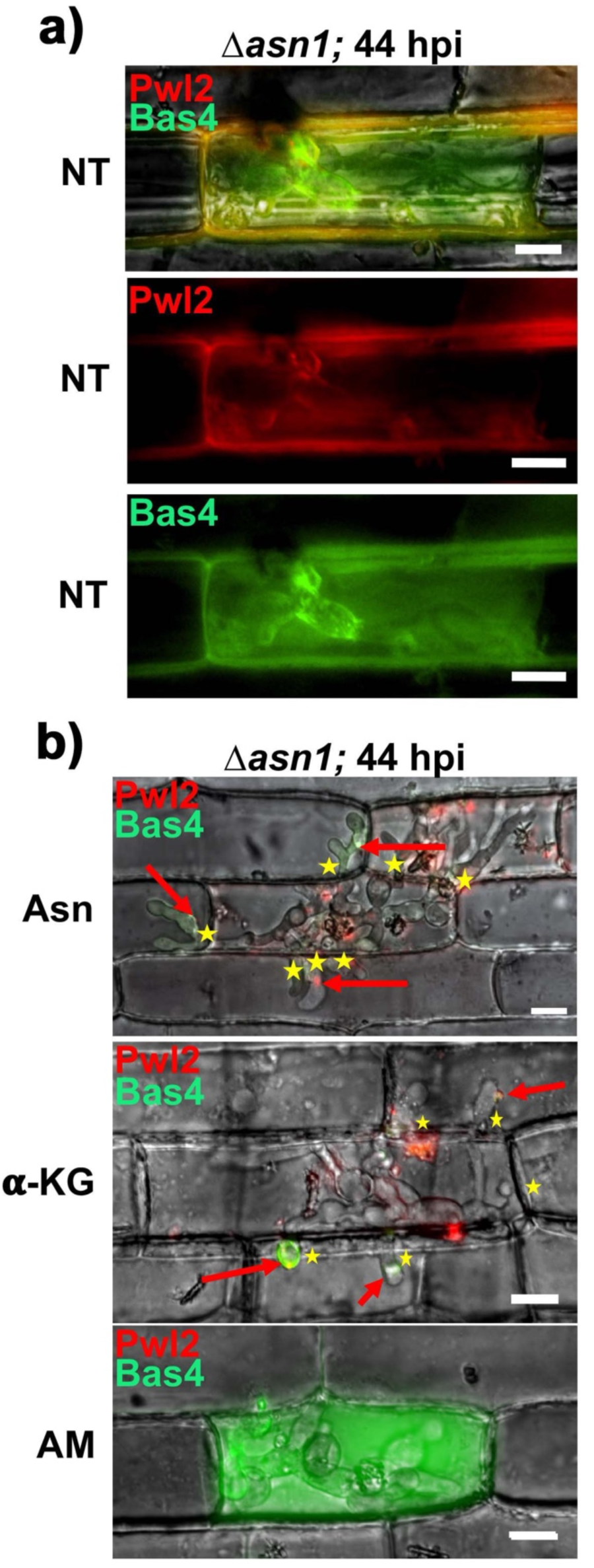
*Δasn1* is required for biotrophic interfacial membrane integrity. **a.** Live cell imaging of detached rice leaf sheaths show how at 44 hpi, an asparagine auxotrophic mutant *Δasn1* expressing Pwl2-mCherry^NLS^ and Bas4-GFP in untreated (NT) cells is abolished for biotrophic growth and leaks Bas4 and Pwl2 into the rice cytoplasm, suggesting biotrophic interfacial membrane integrity is compromised. **b.** Treatment at 36 hpi with 10 mM asparagine or 10 mM *α*-ketoglutarate (as the cell-permable DMKG analogue) promotes *Δasn1* movement to adjacent cells by 44 hpi and remediates biotrophic interfacial integrity, but *Δasn1* is not responsive to 1.5 μM amiodarone hydrochloride treatment. **a,b.** All images are representative of 50 infected rice cells per strain, repeated in triplicate. Asterisks indicate movement of IH into neighbouring cells. Red arrows indicate BICs. Bar is 10 µm.

### *α*-ketoglutarate remediates IH growth following autophagy blocking by 3-MA

Although *α*-ketoglutarate suppresses autophagy, when considering that autophagy in IH is cyclical, and both that *α*-ketoglutarate remediates IH growth following GS-GOGAT inhibition and under amino acid auxotrophy, we hypothesized that *α*-ketoglutarate is an autophagy-derived signaling metabolite. To test this, we asked whether *α*-ketoglutarate treatment could remediate IH growth following autophagy blocking by 3-MA. **Fig 10** shows that it can. Treating WT infected rice leaf sheaths with 3-MA at 36 hpi followed by the replacement of this treatment with water at 40 hpi led, by 44 hpi, to compromised IH growth, the loss of biotrophic interfacial integrity, and the elicitation of robust plant defense responses, compared to the untreated control. However, replacing 3-MA at 40 hpi with *α*-ketoglutarate (as DMKG) resulted, by 44 hpi, in the remediation of IH growth and biotrophic membrane integrity and the suppression of host plant defense responses (**Fig 10**). Our interpretation of these results is that *α*-ketoglutarate by-passes the autophagy block by providing a trigger for TOR reactivation resulting in biotrophic growth. Thus, *α*-ketoglutarate is a likely autophagy-derived signaling metabolite.

**Figure 10.**
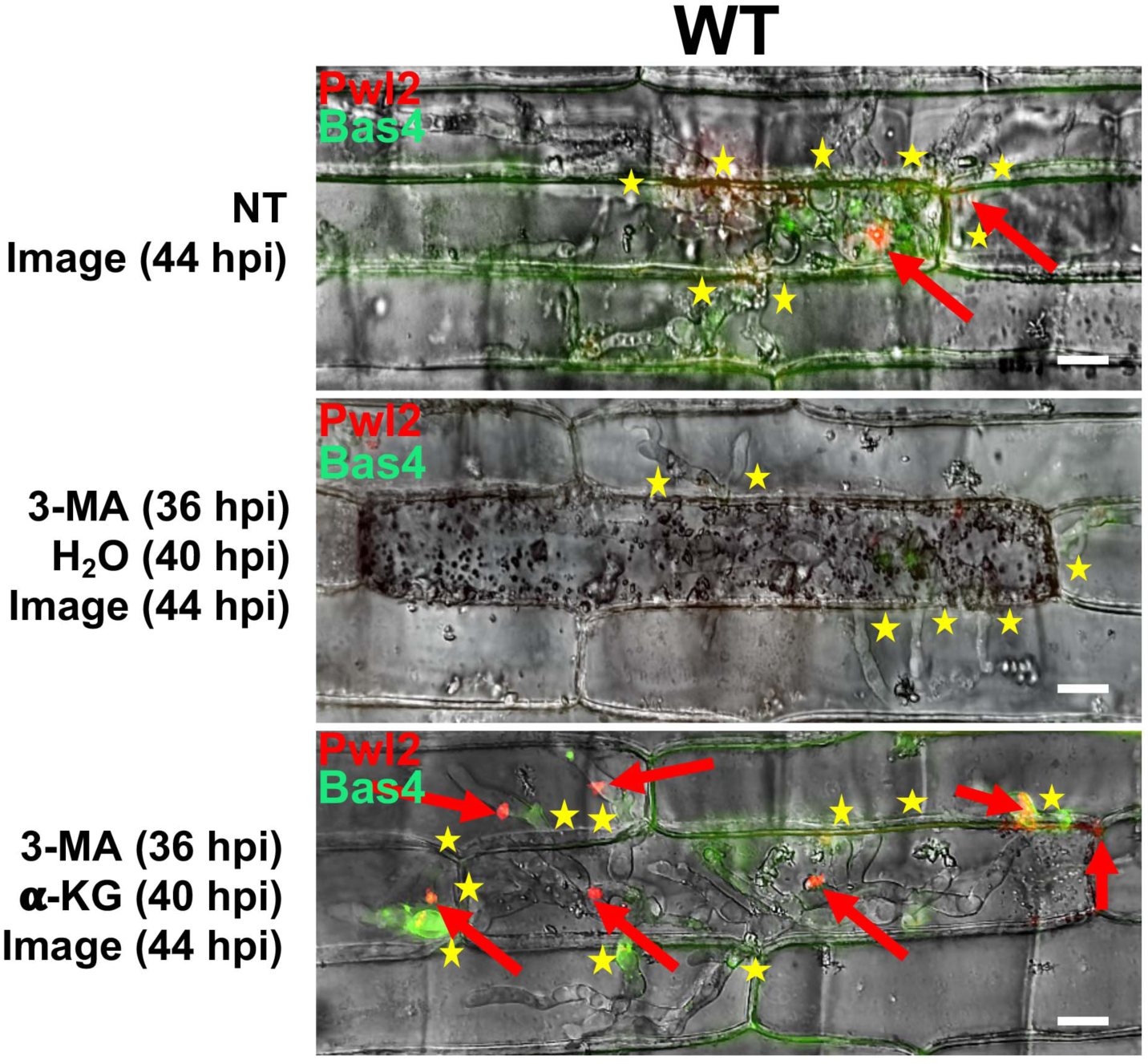
*α*-ketoglutarate treatment remediates IH growth following autophagy blocking by 3-MA. Live cell imaging of detached rice leaf sheaths infected with WT shows how *α*-ketoglutarate can remediate IH growth following autophagy blocking by 3-MA. Infected rice cells were untreated (NT) or treated with 10 mM 3-methyladenine (3-MA) at 36 hpi before this treatment was removed at 40 hpi and replaced with either water or 10 mM *α*-ketoglutarate (as the cell-permable DMKG analogue), as indicated, then imaged at 44 hpi. All images are representative of 50 infected rice cells per strain, repeated in triplicate. Asterisks indicate movement of IH into neighbouring cells. Red arrows indicate BICs. Bar is 10 µm. Merged channel is shown.

### Glutamine treatement remediates *Δimp1* biotrophic growth

Live-cell imaging showed that 10 mM glutamine treatment of detached rice leaf sheaths at 36 hpi also remediated *Δimp1* biotrophic growth by 44 hpi (**Fig S7**). Glutamine was applied here rather than *α*-ketoglutarate (as cell-permeable DMKG) in order to determine if *Δimp1* IH could actively transport amino acids into cells. Because exogenous glutamine treatment remediated *Δimp1* IH growth and cell-to-cell spread, this provides evidence that, like for *Δrim15*, *Δimp1* cannot access sufficient quantities of host glutamine to promote biotrophic growth. These results also point to Imp1 acting upstream of the *α*-ketoglutarate signal.

## Discussion

Many important eukaryotic filamentous plant pathogens exhibit a symptomless biotrophic stage, where microbial cells grow in living host plant cells separated from host cytoplasm by extensive biotrophic interfaces, but little is known about the molecular and metabolic drivers involved. The major findings of this work help to fill these knowledge gaps. We have shown that fungal autophagy is cyclical during growth in host rice cells, and that *α*-ketoglutarate liberated by autophagy reactivates TOR to stimulate biotrophic growth. Moreover, because *α*-ketoglutarate is generated in cells from excess amino acids via glutaminolysis and the GS-GOGAT cycle – processes that are shown here to be essential for biotrophic growth – we provide evidence that *α*-ketoglutarate is an amino acid-sufficiency signal for triggering TOR. Thus, the significance of our work lies in uncovering the metabolic strategies underpinning host infection by a devastating plant pathogen, thereby shedding considerable new light on understanding how the cells of one organism can grow in the cells of another.

All our data together fit the model in **Fig 11**, which highlights the significance of autophagic cycling and *α*-ketoglutarate production in integrating fungal biotrophic growth with interfacial membrane integrity and host defense suppression during growth in living host rice cells. After an initial period of IH elaboration, which does not require *RIM15* but which we have shown previously does require glucose-mediated TOR activation following host penetration (Fernandez et al. 2014), and after initial biotrophic interface construction (which also does not require *RIM15*), further extensive biotrophic growth and interface maintenance is driven by successive cycles of *RIM15-*dependent autophagic flux and *α*-ketoglutarate-mediated TOR reactivation. Autophagy induction requires Imp1, which facilitates membrane homeostasis (Sun et al. 2018), and autophagy induction is blocked by the autophagy inhibitor 3-MA. Rim15 is not required for autophagy induction but, like in yeast, is required for autophagic flux downstream of Imp1. Rim15 also ensures the production of *α*-ketoglutarate from autophagy-liberated amino acids via glutaminolysis and the GS-GOGAT pathway. In this context, in WT, *α*-ketoglutarate acts as an endogenous amino acid sufficiency signal to trigger TOR reactivation, leading to autophagy suppression and IH growth. As excess amino acids are consumed for growth, *α*-ketoglutarate is depleted, TOR is inactivated and autophagy is induced. Discovering that autophagy mediates host-pathogen interactions through the supply of *α*-ketoglutarate is significant because although the function of autophagy in cells is well understood, few physiological roles – including how autophagy-derived metabolites are used by cells – are known (May et al. 2020). Such pivotal roles for autophagy and *α*-ketoglutarate in pathogenicity could be exploited as targets for broad range disease mitigation strategies.

**Figure 11.**
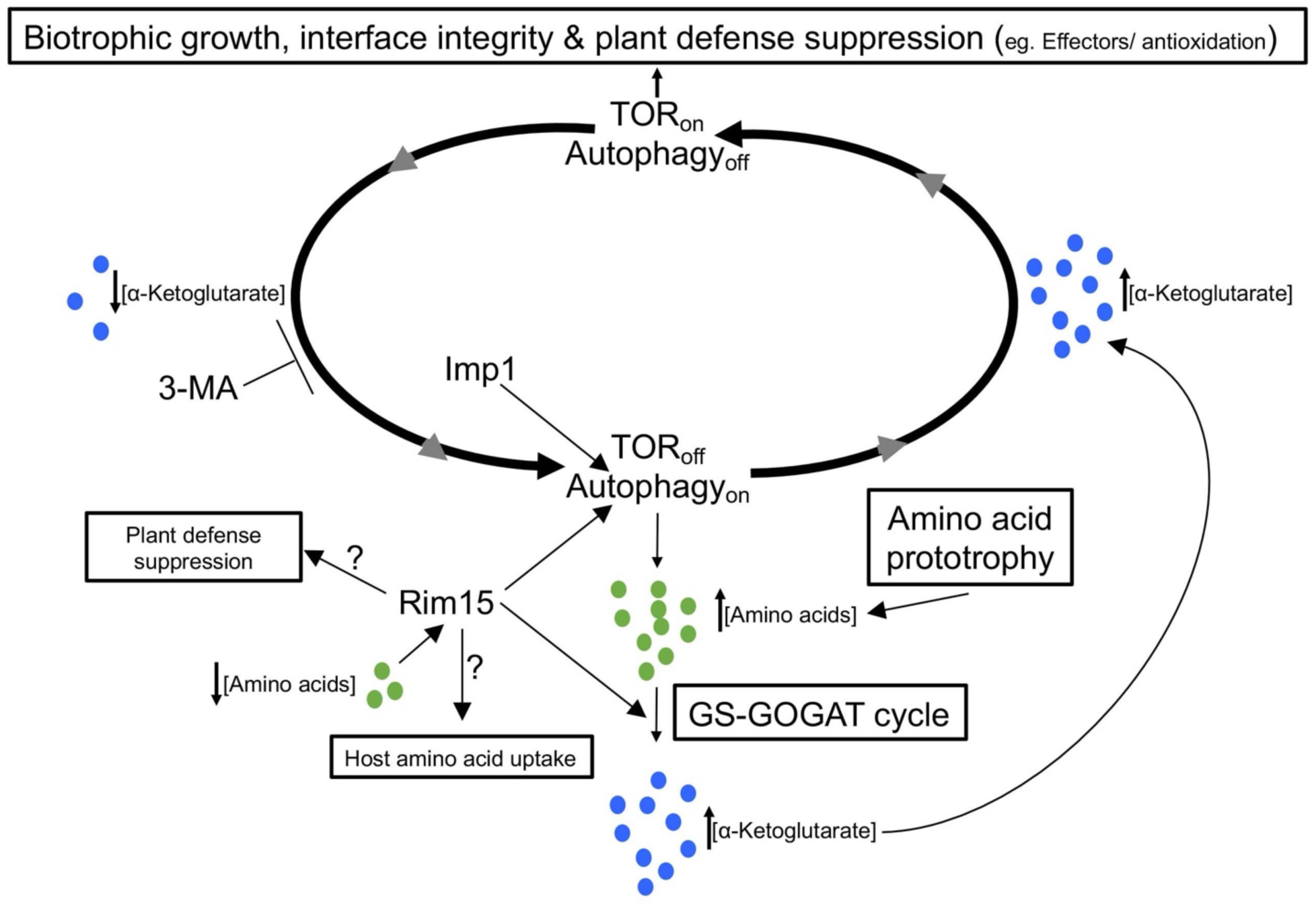
Cycles of autophagy-dependent TOR reactivation mediated by *α*-ketoglutarate drive fungal growth in living host rice cells. Model shows how Rim15-dependent autophagy activity and the GS-GOGAT cycle modulate *α*-ketoglutarate levels in an amino acid prototrophy-dependent manner. Abundant *α*-ketoglutarate is an amino acid sufficiency signal to trigger TOR reactivation and drive biotrophic growth. The role of Imp1 is indicated, and additional roles for Rim15 are postulated. See text for details.

That *α*-ketoglutarate from glutamine and glutamate is the TOR reactivation signal, and not glutamine or gutamate from *α*-ketoglutarate, was suggested by several pieces of evidence, including the remediation of AZS treatment by *α*-ketoglutarate, the observation that *α*-ketoglutarate alone was sufficient to suppress autophagy in vegetative hyphae in the absence of any other nutrients including a nitrogen source, and by the remediation of IH growth by *α*-ketoglutarate following autophagy blocking by 3-MA treatment. The signaling role deduced here for *α*-ketoglutarate has not been previously described for any host-pathogen interactions, but our findings are inline with studies showing *α*-ketoglutarate production from glutaminolysis, and treatment with the cell-permable *α*-ketoglutarate analogue, stimulated mTORC1 activation in human cells (Durán et al. 2012). Moreover, similar to our observations that *α*-ketoglutarate alone can suppress autophagy via TOR reactivation, supplementation with glutamine alone was sufficient to restore mTORC1 activity in mouse embryonic fibroblasts (MEFs) following a prolonged period of amino acid starvation, and mTORC1 reactivation after starvation in MEFs was autophagy-dependent (Tan et al. 2017). Thus, the general principles of the *α*-ketoglutarate-mediated TOR reactivation mechanism we describe here, essential for regulating cell growth under nutrient-limited conditions, might be widespread.

We showed how autophagy-dependent TOR reactivation is dependent on amino acid prototrophy, likely in order to maintain a sufficient and homeostatic pool of amino acids balancing glutamine and glutamate conversion to *α*-ketoglutarate. Treatments suggested that this pool seemingly collapses in *Δasn1* IH, which requires exogenous asparagine to remediate growth. Also, unlike for *Δrim15*, autophagy induction by AM treatment could not compensate for the loss of asparagine from the endogenous amino acid pool. Coupled with the amino acid treatments of *Δrim15*-infected rice leaf sheaths and the effect of *α*-ketoglutarate treatment on suppressing WT autophagy, these results provide the most compelling evidence to date that host nutrient acquisition by *M. oryzae* is very low or possibly, at least for some metabolites, non-existent, perhaps as a means to avoid detection by the host plant. Thus, caution must be taken when considering mitigation strategies based around limiting presumed fungal access to host nutients. However, some limited host nutrient acquisition must occur to facilitate IH growth following TOR reactivation, and a clue as to what this might be is found in the RNAseq data (**Table S1**), where a putative glutamate symporter, MGG_07639, annotated at NCBI as encoding a homologue of excitatory amino acid transporter 2 (EAAT2), is downregulated 10-fold in *Δrim15*-infected rice cells at 36 hpi compared to WT. The significance of this transporter to infection and *M. oryzae* glutamate metabolism will be addressed in the future, but its downregulation in *Δrim15* suggests that at least some host nutrient acquisition occurs *in planta* in a Rim15-dependent manner.

How does Rim15 control autophagic flux and optimize the GS-GOGAT cycle? In yeast, Rim15 can regulate *ATG* gene expression by phosphorylation inhibition of Ume6, a transcriptional repressor of *ATG* genes (Bartholomew et al. 2012; Kim et al. 2021). MGG_08974 is the top hit when interrogating the *M. oryzae* genome at EnsemblFungi with the ScUme6 protein sequence. However, MGG_08974 was not detected in our RNAseq, proteomic or quantitative phosphoproteomic data (**Table S1-S3**). Instead, we noted that phosphorylation levels of the protein product of a *GCN4* homologue (MGG_00602) (known as *cpc-1* in *Neuorospora crassa*) was decreased 4.5 fold in *Δrim15* compared to WT under glutaminolytic growth conditions (**Table S3**), suggesting that the basic leucine zipper (bZIP) transcriptional activator Gcn4 might be a direct or downstream target of Rim15. In yeast, Gcn4 is a master activator of amino acid biosynthetic genes under amino acid starvation conditions, including genes of the GS-GOGAT pathway, and it is also required for nitrogen- and amino acid starvation-induced autophagy gene expression (Natarajan et al. 2001; Tallóczy et al. 2002; Yang et al. 2010; Mittal et al. 2017; Delorme-Axford and Klionsky, 2018; Srinivasan et al. 2020). Thus, in *M. oryzae*, it is conceivable that Rim15-dependent autophagy and Rim15-dependent glutaminolysis processes are both mediated by MoRim15 control of MoGcn4 activity, a relationship not observed in yeast. This putative Rim15-Gcn4 connection will be explored further going forward. Also, glutamate dehydrogenase (Gdh1, MGG_08074) was reduced 10- fold in abundance in *Δrim15* compared to WT under glutaminolytic growth conditions, and phosphorylation levels were reduced 100-fold. Although these reductions might be indirectly due to the loss of Rim15, future work will examine whether a direct interaction betwen Rim15 and Gdh1 occurs in order to stabilize Gdh1 through phosphorylation during glutaminolysis, thus providing an additional fine level of GS-GOGAT cycle control.

TOR reactivation by *α*-ketoglutarate is sufficient to suppress host defenses, but does Rim15 have a TOR-independent role in plant innate immunity suppression? The observation that *Δrim15* induces stronger plant defense responses by 44 hpi than *Δimp1*-infected cells suggests that the loss of autophagy induction and biotrophic interface integrity is not sufficient by itself to trigger such strong plant defenses in *RIM15^+^* strains. If Rim15 has an independent role in plant defense suppression, it is likely to be at the level of ROS detoxification, for several reasons. First, glutathione, a tripeptide comprising glutamate, cysteine and glycine, is required in *M. oryzae* for suppressing host ROS (Rocha and Wilson, 2020) and is reduced in abundance in *Δrim15* vegetative mycelia compared to WT (**Table S5**), likely due to the perturbations in glutamate allocation through the GS-GOGAT cycle. Second, in yeast, Rim15 controls the expression of stress-responsive genes via the transcription factors Msn2/4 and Gis1, including genes required for oxidative stress resistance (Deprez et al. 2018). Although we could not find *M. oryzae MSN2/4* or *GIS1* homologues in our genome-wide data, the protein encoded at MGG_02408, a homologue of yeast Asg1, is 100-fold reduced in phosphorylation levels in *Δrim15* vegetative mycelia compared to WT under glutaminolytic conditions (**Table S3**). Asg1 in yeast is a transcriptional activator of stress response genes, and MoAsg1 might be a direct or downstream target of MoRim15 with a role in neutralizing ROS and maintaining redox balance. Finally, in yeast, Rim15 mediates chronological aging via the modulation of mitochondrial function to reduce endogenous ROS in the stationary phase (Deprez et al. 2018). The loss of a similar mitochondial integrity function in *M. oryzae Δrim15* strains would impair cellular redox balance, resulting in the elevated host ROS observed at 44 hpi. These potential oxidative stress-related and ROS detoxification roles of *M. oryzae* Rim15 will be investigated in the future.

To conclude, this work provides an unparalleled description of the core metabolic strategies required for establishing fungal biotrophic growth in host plant cells, and points to Rim15-mediated networks of gene, protein and metabolic interactions underlying infection. As the climate changes, the number and assemblages of eukaryotic filamentous plant pathogens pressuring food security will change and new threats will emerge (Chaloner et al. 2021). Combatting these threats in a robust and sustainable manner will require a more detailed and comprehensive understanding of the molecular factors facilitating pathogen invasive growth in host plant cells. By deducing how *α*-ketoglutarate, revealed here as a key integrator of carbon and nitrogen metabolism, connects autophagic cycling and TOR signaling to support pathogen invasive growth and metabolic homeostasis in the absence of extensive nutrient acquisition from the host, we provide a solid foundation for future studies probing and interpreting fungal metabolism in host cells. Such studies might foster the discovery of new and targetable pathogen weaknesses with broad applicability.

## Materials and methods

### Fungal strains and growth conditions

The *M. oryzae* Guy11 rice isolate was used as the wild type (WT) strain for this study (Wilson and Talbot, 2009). Strains used in this study are listed in **Table S6** and are maintained as filter stocks in the Wilson lab. Strains were grown at 26 °C on solid and liquid CM or minimal media as described in detail previously (Li et al. 2015; Sun et al. 2018). *In vitro* fungal physiological analyses were performed as described previously (Sun et al. 2018), and specific details are provided in the text.

### Plant inoculation assays and live-cell imaging

For whole plant inoculations, spores of the indicated strains were harvested from colonies grown on solid oatmeal agar plates for 14 days and applied to 3-week-old rice seedlings of the susceptible cultivar CO- 39 at a rate of 1 ×10^5^ spores ml^−1^. Images were recorded at 5 days post inoculation. For live-cell imaging, spores of the indicated strains were harvested from 14-day-old colonies grown on solid oatmeal agar plates and inoculated at a rate of 2 ×10^4^ spores ml^−1^ onto detached rice leaf sheaths from 4-week-old rice seedlings of the susceptible cultivar CO-39, as described previously (Sun et al. 2018). Leaf sheath cells were imaged at the indicated times using a Nikon Eclipse Ni-E upright microscope and NIS Elements software. Unless otherwise specified, treatments were added at 36 hpi and viewed at 44 hpi. Treatments were added at the concentrations indicated. Treatments were purchased from Sigma-Aldrich, USA. Cells were stained with 1 mg ml^-1^ 3,3′-diaminobenzidine (DAB, Sigma-Aldrich) as previously described (Marroquin-Guzman et al. 2017). All treatment assays were performed at least in triplicate and representative images are shown.

Detached rice leaf sheath assays and artificial hydrophobic surfaces were used to quantify appressorium formation rates. Appressorium penetration rates and rates of IH cell-to-cell movement to adjacent cells were determined using detached rice leaf sheaths. Appressorial formation rates were determined by examining how many of 50 spores had formed appressoria by 24 hpi on artificial hydrophobic surfaces (after applying at a rate of 1 x 10^4^ spores ml^-1^) or on rice leaf sheath surfaces, repeated in triplicate. Penetration rates were determined by examining how many of 50 appressoria on one rice leaf sheath surface had penetrated into underlying cells by 30 hpi, repeated in triplicate. Rates of cell-to-cell movement were determined by observing how many of 50 primary infected rice cells had formed IH in adjacent cells by 48 hpi, repeated in triplicate.

### Generation of targeted gene deletion mutant strains in *M. oryzae*

Genes were disrupted in *M. oryzae* using the split marker technique (Sun et al. 2018) and the primers are listed in **Table S7**. *RIM15* and *ASN1* genes were replaced with the *ILV2* gene conferring resistance to sulphonyl urea. Transformants were confirmed by PCR. At least two independent deletants per gene knockout were characterized fully and shown to be indistinguishable.

### Generation of strains expressing fluorescently labelled proteins

*M. oryzae* gene sequences obtained from the *M. oryzae* genome database (http://fungi.ensembl.org/Magnaporthe_oryzae/Info/Index) were used to design the primers in **Table S7** that were used for vector construction. The full length CDS of *ATG8* was amplified from the Guy11 strain with RP27-NGFP-ATG8-F /-R primers. The end of the PCR product contained 15∼20 bases homologous to the pGTN vector. Purified PCR product was fused with linearized pGTN, which was digested with Not I using the In-Fusion HD enzyme kit. The reaction mixture was transferred into the *E. coli* DH 5a strain with ampicillin antibiotic screening, and all colonies were identified by PCR and plasmid integrity verified by sequencing. The vector was then transformed into WT and *Δrim15* protoplasts, respectively. All *M. oryzae* transformants were screened for geneticin resistance and confirmed by PCR. A similar procedure was used to construct the *RIM15-GFP* vector, except pGTN was linearized with Hind III and BamH I to insert *RIM15* upstream of *GFP*. To generate *Δrim15* strains expressing Pwl2-mCherry^NLS^ and Bas4-GFP, the *Δrim15* parental strain was transformed with the pBV591 vector (Khang et al. 2010) and selected using hygromycin resistance. To generate *Δasn1* strains expressing Pwl2-mCherry^NLS^ and Bas4-GFP, *ASN1* was disrupted in our *ASN1^+^* strain carrying pBV591. Following transformation, at least 5 independent mutant strains were screened for fluorescence in rice leaf sheath assays.

### RNA extraction and RNA seq analysis

4-week-old rice detached rice leaf sheaths were inoculated with spores of the WT and *Δrim15* mutant strain at a rate of 1×10^5^ spores mL^-1^. Detached rice leaf sheaths inoculated with mock solution (0.2% gelatin) was used as the control. At 36 hpi, the inoculated leaf sheaths were harvested and flash frozen in liquid nitrogen. Three biological replicates of each treatment were used for the following RNA sequencing analysis. For RNA extraction, leaf sheath tissues were ground in liquid nitrogen, and total RNA was extracted using the TRIzol reagent (Invitrogen, US), according to the manufacturer’s standard procedure. Following removal of contaminating genomic DNA using DNase I, total RNA was purified with PureLink RNA Mini kit (Thermo Fisher Scientific, US) by following the manufacture’s instruction manual. Total RNA quality was estimated by running on a denatured agarose gel, and the RNA quantity was measured using a Nanodrop Spectrophotometer. RNA integrity and purity were evaluated using the Agilent Technologies 2100 Bioanalyzer, and high quality (RIN ≥7.0) was confirmed for all samples. For each sample, 3 μg of total RNA at a concentration of more than 100 ng/ul was used in library construction.

For library construction and sequencing, Poly(A) RNA sequencing library was prepared following Illumina’s TruSeq-stranded-mRNA sample preparation protocol. Poly(A) tail-containing mRNAs were isolated from total RNA using oligo-(dT) magnetic beads with two rounds of purification. Then, the poly(A) mRNA fractions were cleaved into smaller fragments using divalent cation buffer in elevated temperature. The cleaved RNA fragments were reverse-transcribed to create the final cDNA library in accordance with a strand-specific library preparation by dUTP method. Quality control analysis and quantification of the sequencing library were performed using Agilent Technologies 2100 Bioanalyzer High Sensitivity DNA Chip. The average insert size for the paired-end libraries was 300 ± 50 bp. Sequencing of generated cDNA libraries was performed with 150-nucleotide pair-end reads on Illumina’s NovaSeq 6000 sequencing system at LC Sciences (US) following the vendor’s recommended protocol. Three biological replications were used for RNA-seq experiments.

For transcriptomic analysis of differentially expressed genes, with reads mapping to the reference genome, raw sequencing data were first filtered by removing contaminated adaptor sequences, low quality bases and undetermined bases using CUTADAPT (Martin et al. 2011) and perl scripts in LC Science. Then, sequence quality was verified using FastQC (http://www.bioinformatics.babraham.ac.uk/projects/fastqc/). Following removal of 9 bp of low quality bases from the 5’-ends of the reads using SEQTK (version 1.2) (https://github.com/lh3/seqtk), valid reads in high quality were mapped to the *Magnaporthe oryzae* genome assembly (https://ftp.ncbi.nlm.nih.gov/genomes/all/GCF/000/002/495/GCF_000002495.2_MG8/) using SUBREAD for Unix (version 2.0) (Liao et al. 2013). The alignment was carried out for read groups of all samples, including all replications of WT, Δ*rim15*, and mock control. Gene annotation as GTF (version 2.2) was obtained from NCBI, corresponding to the MG8 genome assembly GCF_000002495.2. Read counts per-gene were computed using FEATURECOUNTS (Liao et al. 2014) from the SUBREAD (version 2.0). After computing counts per million (CPM), the count data were used in the identification of differentially expressed genes using DESEQ2 (version 1.14.0) (Love et al. 2014) by comparing the expression of genes in samples inoculated with Δ*rim15* mutant, WT, and mock control, with differentially expressed genes defined as having adjusted *P* value < 0.05 and absolute log_2_ of fold change between test and control value of ≥1. Genes that were defined as non-differentially expressed were those having adjusted *P* value ≥0.05 or absolute log_2_ of fold change between test and control value of <1. All bioinformatics analyses following the adaptor removal were performed utilizing the High Performance Computing Resources in Holland Computing Center of the University of Nebraska, USA.

### Protein extraction, proteomics and phosphoprotein enrichment

Strains were grown as mycelia in liquid CM for 42 hr before shifting to minimal media with 0.5 mM glutamine as the sole carbon and nitrogen source for 16 hr.

For the proteomics experiments, 0.3g of wet mycelia per strain (in triplicate) was lysed in 1 mL lysis buffer consisting of 7M urea, 2M thiourea, 5 mM DTT, 100mM tris/HCl pH 7.8, and containing 1X complete EDTA-free protease inhibitor and 1X PhosStop phosphatase inhibitor, for 10 min at 20 Hz on a mechanical tissue lyser. Protein was precipitated with acetone and the pellet was washed and redissolved in the lysis buffer containing 2X PhosStop phosphatase inhibitor. The proteins were assayed using the CBX kit (G-Bioscience). An aliquot of proteins was reduced with DTT and alkylated with iodoacetamide prior to digestion with LysC and then trypsin. Each digest was analyzed by nanoLC-MS/MS using a 2h gradient on a Waters CSH 0.075 mm x 250 mm C18 column feeding into an Orbitrap Eclipse mass spectrometer. However, an unknown component interfered with the chromatography and samples were rerun after offline solid-phase C18 clean-up (Waters SepPak, 100mg 1cc syringe cartridges).

For phosphoprotein enrichment, of the 1mg of wet mycelia from each strain (in triplicate), 0.95mg was used for TiO2 phosphopeptide enrichment with lactic acid. Samples were subjected to offline solid-phase C18 clean-up (Waters SepPak, 100mg 1cc syringe cartridges). Each cleaned sample was then analyzed by nanoLC-MS/MS using a 2h gradient on a Waters CSH 0.075mmx250mm C18 column feeding into an Orbitrap Eclipse mass spectrometer.

Quantification of the proteins and phosphoproteins was performed separately using Proteome Discoverer (Thermo; version 2.4). All MS/MS samples were searched using Mascot (Matrix Science, London, UK; version 2.6.2). Mascot was set up to search the cRAP_20150130.fasta (124 entries); uniprot-refprot_UP000009058_Magnaporthe_oryzae 20210511 (12,791 sequences); assuming the digestion enzyme trypsin. Mascot was searched with a fragment ion mass tolerance of 0.06 Da and a parent ion tolerance of 10.0 PPM. For the proteomics experiment, deamidation of asparagine and glutamine and oxidation of methionine were specified in Mascot as variable modifications while carbamidomethyl of cysteine was specified as fixed modification. For the phosphoproteomics experiment, deamidation of asparagine and glutamine, oxidation of methionine, phosphorylation of serine, threonine and tyrosine, and acetylation of N-term were specified in Mascot as variable modifications while carbamidomethyl of cysteine was specified as fixed modification. Peptides were validated by Percolator with a 0.01 posterior error probability (PEP) threshold. The data were searched using a decoy database to set the false discovery rate to 1% (high confidence). Only proteins with a minimum of 2 peptides and 5 PSMs were reported. For phosphoproteins, the minimum was 1 phosphopeptide and 3 PSMs. The localization probability of the phosphorylation sites is calculated using PtmRS (Taus et al. 2011). Probabilities are indicated in parenthesis next to the amino acid residue. If there is no probability indicated, this means that the phosphorylation of the peptide was not confidently localized.

The peptides and phosphopeptides were quantified using the precursor abundance based on intensity. Normalized and scaled abundances are reported. The peak abundance was normalized using total peptide amount. The peptide group abundances are summed for each sample and the maximum sum for all files is determined. The normalization factor used is the factor of the sum of the sample and the maximum sum in all files. Then, abundances were scaled so that the average of the abundances is equal to 100 for each sample. The imputation mode was used to fill the gaps from missing values. The protein and phosphoprotein ratios are calculated using summed abundance for each replicate separately and the geometric median of the resulting ratios is used as the protein ratios. The significance of differential expression is tested using a t-test which provides a p-value and an adjusted p-value using the Benjamini-Hochberg method for all the calculated ratios.

### Metabolite extraction and Metabolomics

Strains were grown as mycelia in liquid CM for 42 hr before shifting to minimal media with 0.5 mM glutamine as the sole carbon and nitrogen source for 16 hr. Samples were lyophilized before metabolite extraction.

For the metabolomic results in **Table S4**, an aliquot of 20-25 mg of each sample was extracted for polar compounds using the chloroform/methanol/water extraction (Folch Method). The extracts were dried down and derivatized for GCMS using MSTFA+TMCS. The samples were run alongside the mixture of retention index C10-25 alkanes for identification. The data was analyzed using MS-Dial (https://doi.org/10.1038/nmeth.3393) for peak detection, deconvolution, alignment, quantification and identification. The library used was the curated Kovats RI (28,220 compounds, which includes the Fiehn, RIKEN and MoNA databases). The peaks were reviewed and the final list of compounds with RI similarities >90% were reported. Highlighted in red in **Table S4** are compounds with ID confirmed by authentic standards or by complementary search of the NIST14 library using NIST-MS. The peak area was normalized based on the total ion chromatogram for the assigned metabolites using sum normalization in MetaboAnalyst 5.0.

For the metabolomic results in **Table S5**, the samples were suspended in 500 uL of extraction solution on dry ice, and disrupted by 4 cycles of the bullet blender at a setting of “7” for 3 minutes each. The suspension was centrifuged for 10 min at 15,000 xg. The supernatant was transferred into a second tube and evaporated using a speed Vac at 4 °C. The pellet was resuspended into 100 uL of LC-Grade water and transferred into V-vials and placed in the autosampler. The sample concentrations are reported in micromolar units, and the metabolite levels were normalized by weight.

## Acknowledgements

This work was supported by National Science Foundation funding (IOS-1758805) to RAW. ZG was supported by funding from the China Scholarship Council. We thank Dr. Sophie Alvarez of the Center for Biotechnology, University of Nebraska-Lincoln, for the (phospho)proteomic and metabolomic analyses in Tables S2-S4. We thank Dr. Javier Seravalli of the Redox Biology Center, University of Nebraska-Lincoln, for the metabolomic analysis in Table S5. The bioinformatics analysis was completed using the supercomputing resources in Holland Computing Center of the University of Nebraska, USA, which receives support from the Nebraska Research Initiative.

## Author contributions

RAW conceived the project and obtained funding. GL and RAW designed the experiments and interpreted the data. GL, ZG, ND and ROR performed the experiments, specifically, GL generated and characterized the *Δrim15* mutant strains in Guy11, pBV591 and histone H1 parental backgrounds; GL constructed the *RIM15-GFP* vector ; ROR generated the *Δasn1* mutant strain carrying pBV591; ZG characterized *Δasn1* expressing Pwl2-mChery^NLS^ Bas4-GFP, constructed the *GFP-ATG8* vector and generated strains expressing Rim15-GFP and GFP-Atg8; GL and ZG performed all the microscopy; GL generated material for RNAseq analysis and performed all bioinformatic analyses; ND generated material for proteomic, phosphoproteomic and metabolomic analyses. GL composed the figure panels with input from ZG. RAW wrote the manuscript and finalized the figures, with contributions from the other authors.

---

**Supplemental Figure 1.**
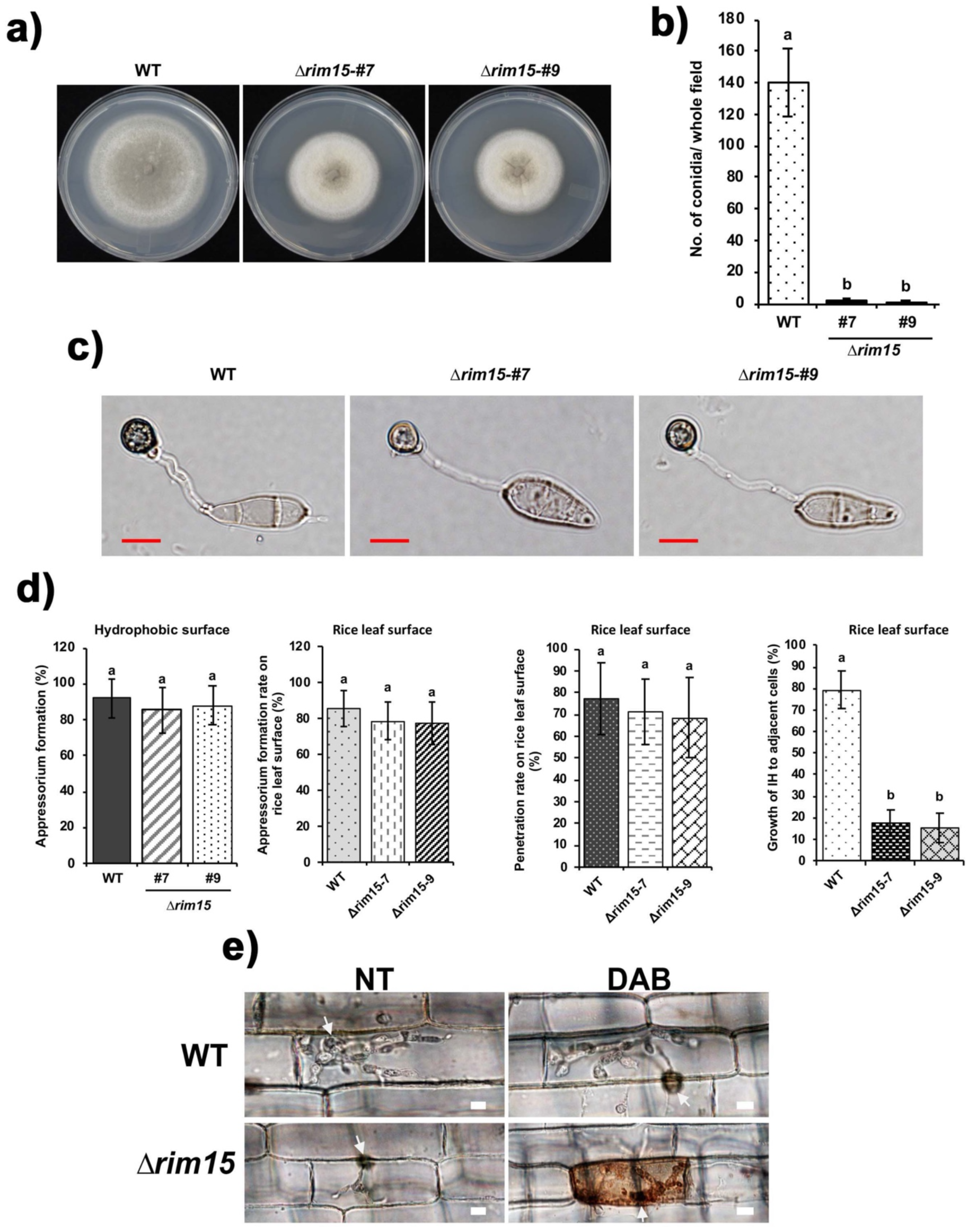
R*I*M15 is required for IH cell-to-cell growth and for suppressing the host ROS burst. **a.** Plate test showing how on complete media (CM), the radial growth of two independent *Δrim15* gene knockout mutants was reduced compared to WT. **b.** Sporulation rates of two independent *Δrim15* mutants were reduced on CM compared to WT. Spores of the indicated strains were harvested from CM at 10 days, and the number of spores was counted using a haemocytometer under a dissecting microscope. Results are the mean of three independent measurements. Error bars indicate standard deviation. For each column, different letters indicate significant differences at *P* ≤ 0.05 using the Student-Newman-Keuls test. **c.** Loss of *RIM15* did not impair appressorium formation on artificial hydrophobic surfaces. Spores were harvested from 14-day-old colonies of WT and two Δ*rim15* mutant strains growing on oatmeal agar medium and then resuspended at 2×10^4^ spores ml^-1^. 200 μl of the spore suspension was inoculated onto a hydrophobic plastic cover slip positioned in a humid compartment and then incubated at 25°C for 24 hr in the dark. Scale bar is 10 μm. Experiments were repeated in triplicate. **d.** Graphs showing how appressorium formation rates on artificial hydrophobic surfaces and on rice leaf surfaces at 24 hpi, and appressorial penetration rates on rice leaf surfaces at 30 hpi, were not significantly (P ≤ 0.05) different between *Δrim15* mutant strains and WT. However, cell-to-cell movement rates at 48 hpi were significantly (P ≤ 0.05) reduced in two independents *Δrim15* mutant strains compared to WT. For appressorium formation and penetration rates, 50 spores or appressoria, respectively, were observed for each strain, repeated in triplicate. For IH movement rates, 50 primary infected cells were observed for each strain. Values are the means of three replicates. Bars are standard deviation. Bars with the same letters are not significantly different (Student’s t-test, P≤0.05). **e.** Live cell-imaging of detached rice leaf sheaths infected with the indicated strains shows how loss of *RIM15* elicits an oxidative burst in *Δrim15*- infected host rice cells. Leaf sheaths were stained with 3,3′-Diaminobenzidine (DAB) and imaged at 36 hpi. White arrows indicate appressorial penetration sites. All images are representative of 50 infected rice cells per strain, repeated in triplicate. Scale bar is 10 µm.

**Supplemental Figure 2.**
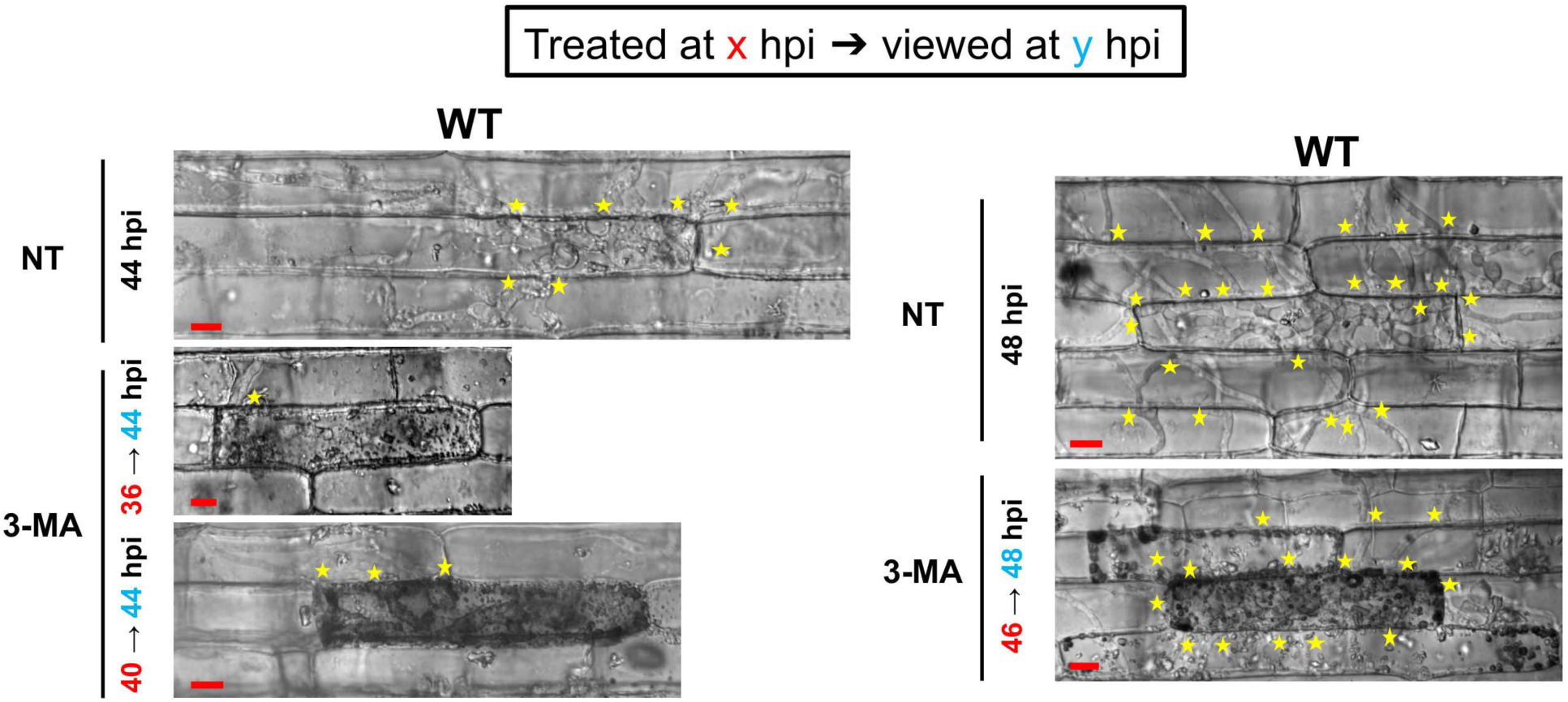
Inhibiting autophagic cycling during biotrophy abolishes IH growth. Live-cell imaging of WT-infected detached rice leaf sheaths shows that, when viewed at the indicated times following 3-methlyadenine (3-MA) treatment, compared to the untreated controls, inhibiting autophagy prevented further IH growth. Asterisks indicate movement of IH into neighbouring cells. NT is no treatment. Bar is 10 µm.

**Supplemental Figure 3.**
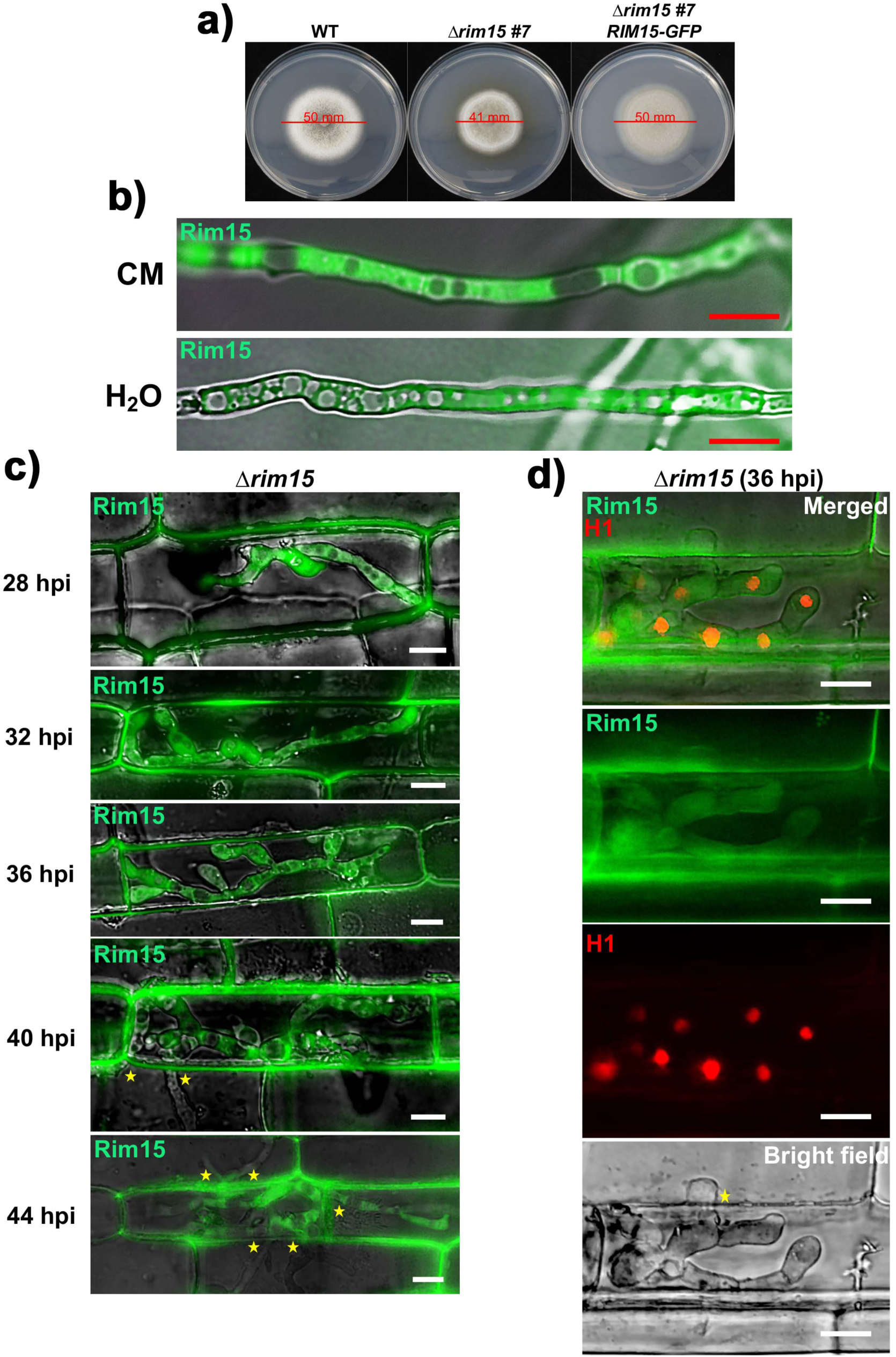
Rim15 localizes to hyphal cytoplasm. **a.** Complementation of *Δrim15* with *RIM15-GFP* restored radial growth on CM media, thus *RIM15-GFP* is functional in *Δrim15*. **b.** Micrograph shows that in vegetative hyphae, Rim15-GFP remains cytoplasmic under nutrient-rich (CM) and nutrient-starvation (water) culture shake conditions. The *Δrim15 RIM15-GFP* complementation strain was grown in liquid CM for 42 hr. After washing with water, vegetative hyphae were transferred into fresh liquid CM or water for a further 3.5 hr before imaging. Bar is 10 µm. Merged channel is shown. **c.** Live-cell imaging at the indicated times of detached rice leaf sheaths infected with the *Δrim15 RIM15-GFP* complementation strain shows that Rim15-GFP localizes to IH cytoplasm throughout biotrophy. Asterisks indicate movement of IH into neighbouring cells. Bar is 10 µm. Merged channel is shown. **d.** Live-cell imaging at 36 hpi of detached rice leaf sheaths infected with a *RIM15^+^* strain expressing histone H1-RFP and Rim15-GFP shows that Rim15-GFP does not co-localize with the nucleus. Asterisks indicate movement of IH into neighbouring cells. Bar is 10 µm.

**Supplemental Figure 4.**
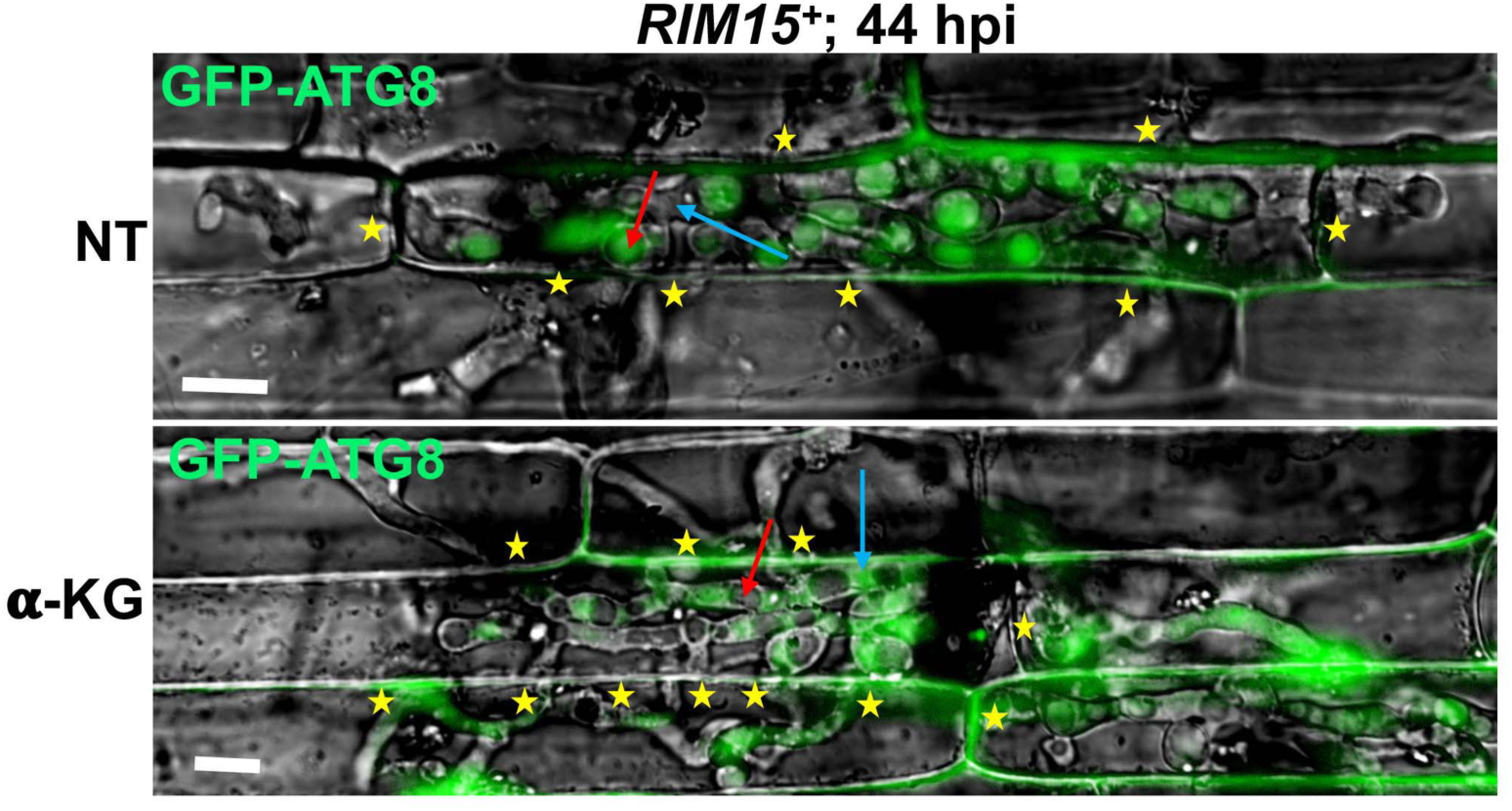
*α*-ketoglutarate treatment suppresses autophagy in WT at 44 hpi. Live-imaging of detached rice leaf sheaths infected with a *RIM15^+^* strain expressing GFP-Atg8 shows that *α*-ketoglutarate treatment (as the cell-permeable DMKG analog) at 36 hpi had, by 44 hpi, suppressed autophagic activity compared to the untreated (NT) control. Examples of vacuoles are indicated with red arrows, examples of cytoplasm are indicated with blue arrows. Asterisks indicate movement of IH into neighbouring cells. Bar is 10 µm. Merged channel is shown.

**Supplemental Figure 5.**
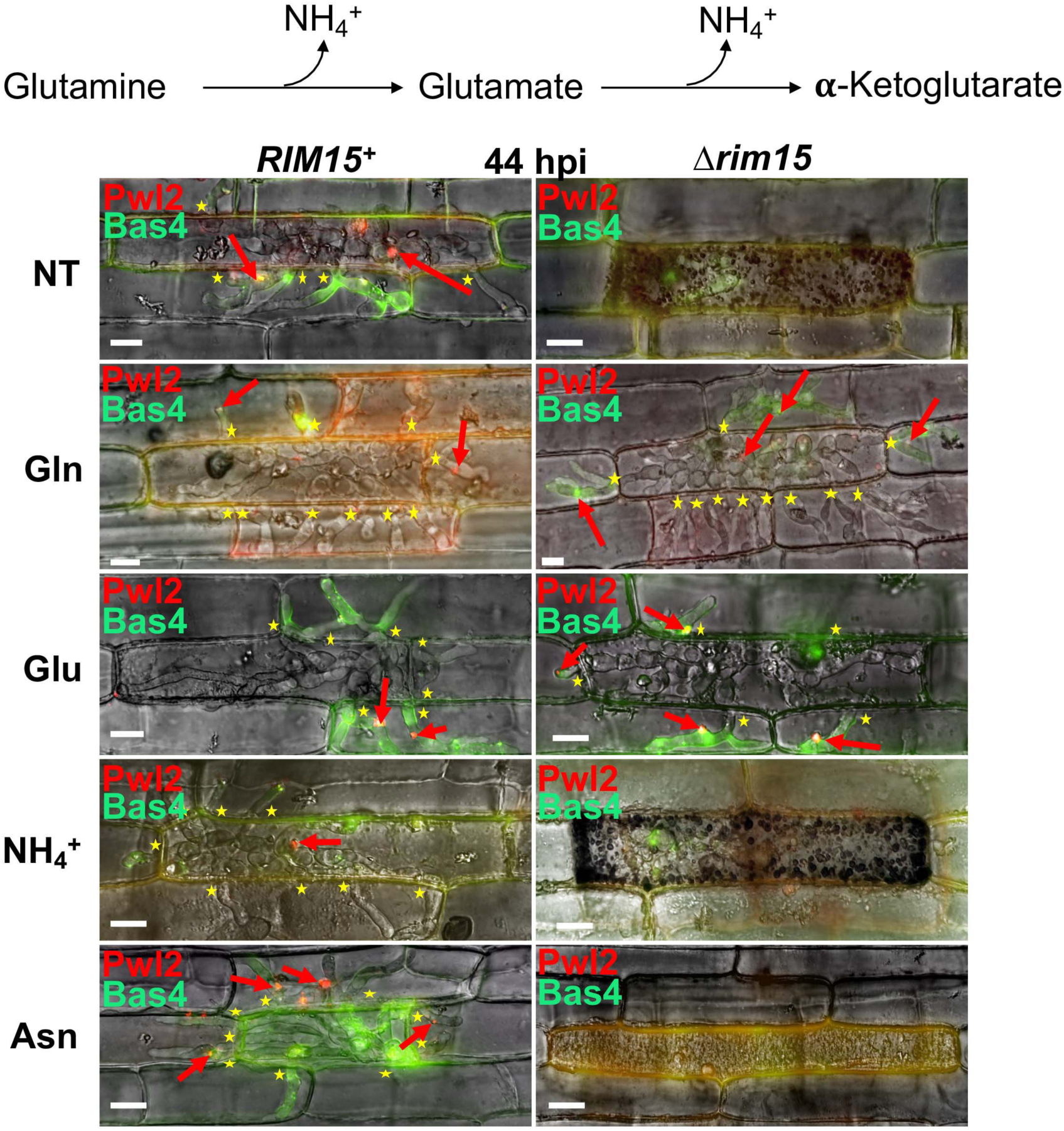
Glutamine and glutamate but not ammonium or asparagine treatments remediated *Δrim15* IH growth and biotrophic interface membrane integrity. Live-cell imaging of detached rice leaf sheaths infected with the indicated strains expressing Pwl2-mCherry^NLS^ and Bas4-GFP show that treatment at 36 hpi with 10 mM of the glutaminolytic amino acids glutamine and glutamate (as L-Glutamic acid monosodium salt hydrate), but not 10 mM ammonium tartrate (NH_4_^+^) or 10 mM asparagine, remediated *Δrim15* biotrophic growth in host rice cells by 44 hpi. Red arrows indicate BICs. Asterisks indicate movement of IH into neighbouring cells. Bar is 10 µm. Merged channel is shown. NT is no treatment.

**Supplemental Figure 6.**
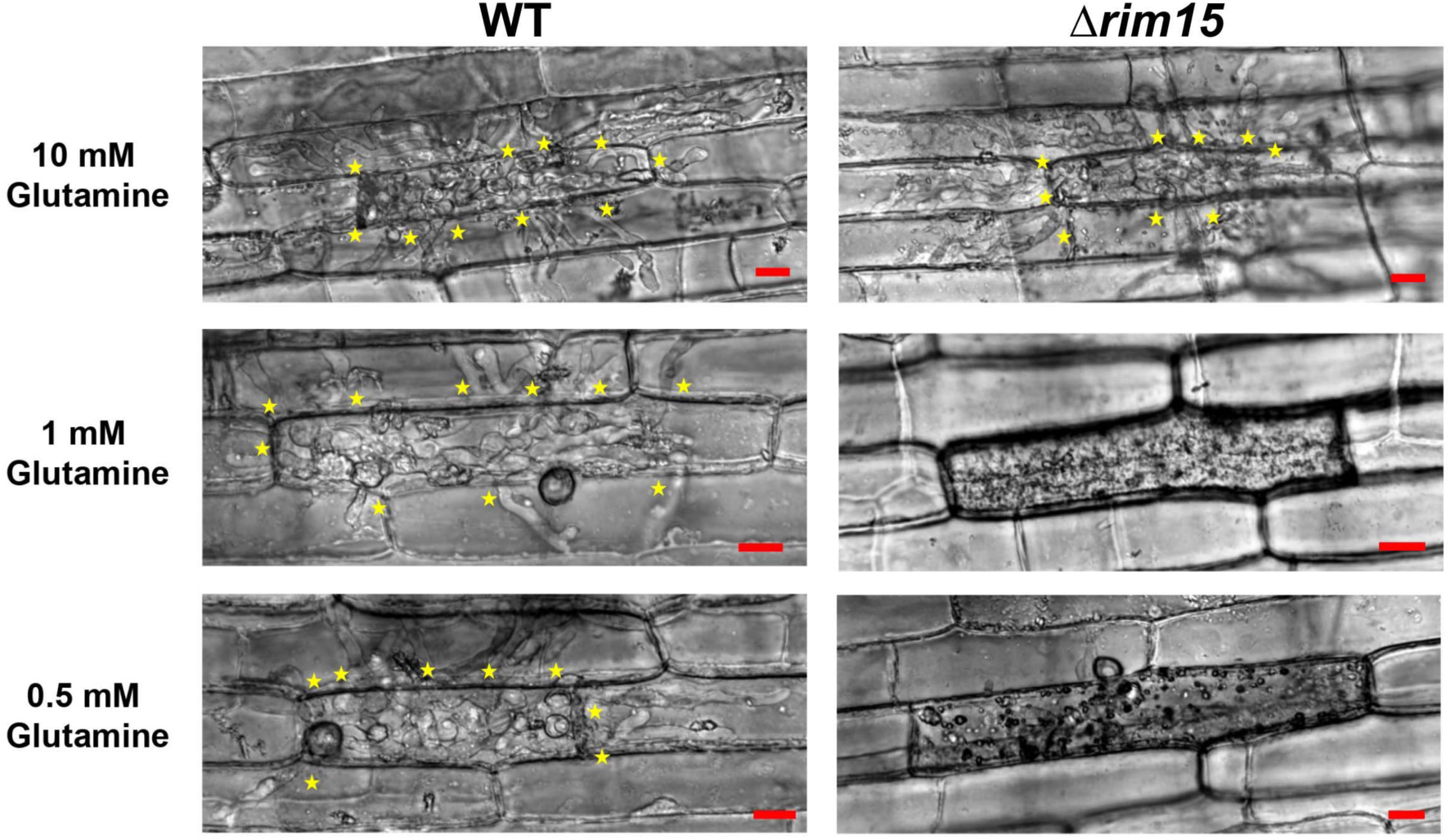
Remediation of *Δrim15* biotrophic growth by glutamine is concentration dependent. Live-cell imaging at 44 hpi of detached rice leaf sheaths infected with the indicated strains shows that glutamine treatment at 36 hpi did not remediate *Δrim15* biotrophic growth when added at low concentrations. Asterisks indicate movement of IH into neighbouring cells. Bar is 10 µm. Bright field channel is shown.

**Supplemental Figure 7.**
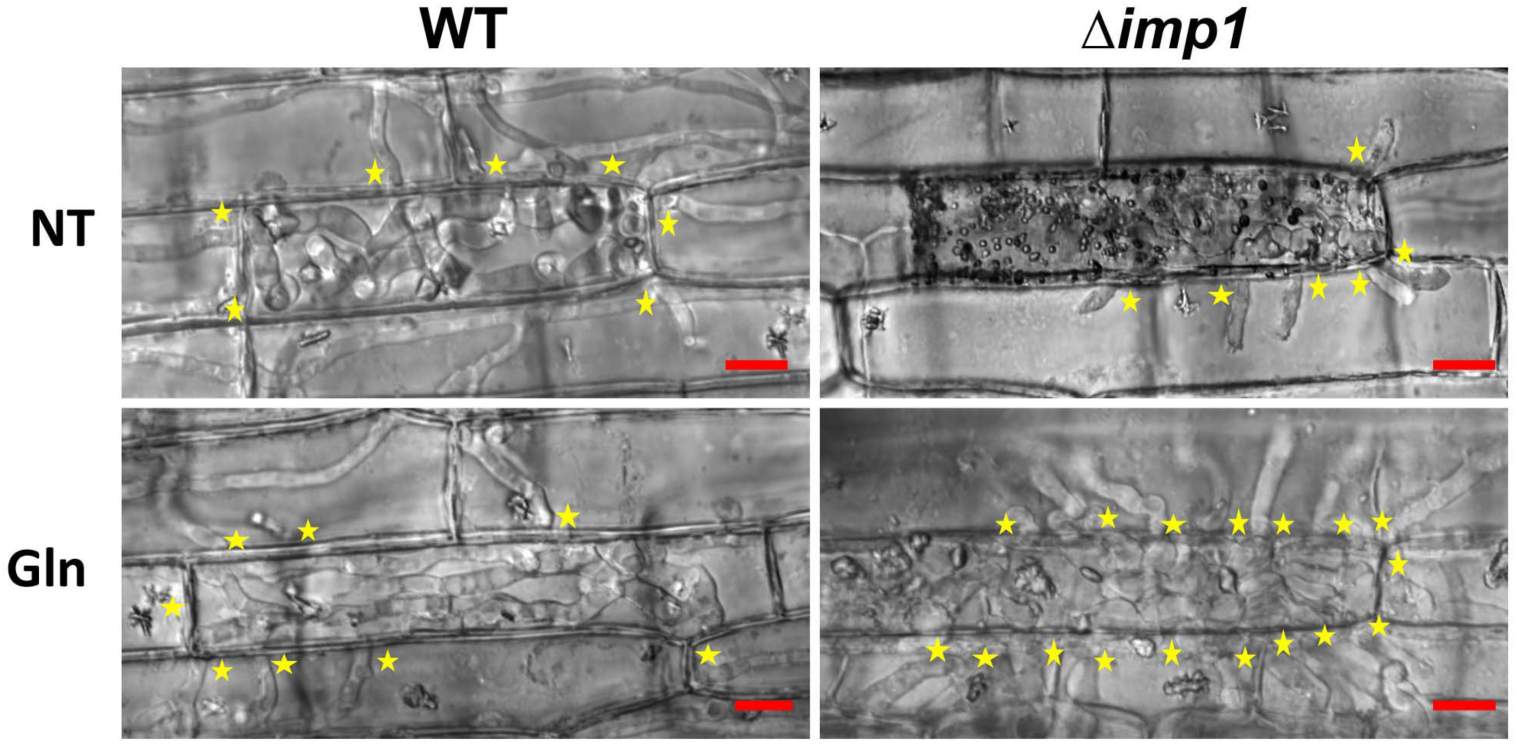
Glutamine treatment remediates *Δimp1* IH growth. Live-cell imaging at 44 hpi of detached rice leaf sheaths infected with the indicated strains shows that 10 mM glutamine treatment at 36 hpi remediated *Δimp1* biotrophic growth. Asterisks indicate movement of IH into neighbouring cells. Bar is 10 µm. Bright field channel is shown. NT is no treatment.

## References

Bartholomew CR, Suzuki T, Du Z, Backues SK, Jin M, Lynch-Day MA, Umekawa M, Kamath A, Zhao M, Xie Z, Inoki K, Klionsky DJ (2012) Ume6 transcription factor is part of a signaling cascade that regulates autophagy. Proc Natl Acad Sci U S A 109: 11206–11210. doi: 10.1073/pnas.1200313109

Chaloner, TM, Gurr, SJ, Bebber, DP (2021) Plant pathogen infection risk tracks global crop yields under climate change. Nat Clim Chang 11, 710–715. doi.org/10.1038/s41558-021-01104-8

Dean RA, Talbot NJ, Ebbole DJ, Farman ML, Mitchell TK, Orbach MJ, Thon M, Kulkarni R, Xu JR, Pan H, Read ND, Lee YH, Carbone I, Brown D, Oh YY, Donofrio N, Jeong JS, Soanes DM, Djonovic S, Kolomiets E, Rehmeyer C, Li W, Harding M, Kim S, Lebrun MH, Bohnert H, Coughlan S, Butler J, Calvo S, Ma LJ, Nicol R, Purcell S, Nusbaum C, Galagan JE, Birren BW (2005) The genome sequence of the rice blast fungus *Magnaporthe grisea*. Nature 434: 980–986. doi: 10.1038/nature03449

Delorme-Axford E, Klionsky DJ (2018) Transcriptional and post-transcriptional regulation of autophagy in the yeast *Saccharomyces cerevisiae*. J Biol Chem 293: 5396–5403. doi: 10.1074/jbc.R117.804641

Deprez MA, Eskes E, Winderickx J, Wilms T (2018) The TORC1-Sch9 pathway as a crucial mediator of chronological lifespan in the yeast *Saccharomyces cerevisiae*. FEMS Yeast Res 18(5). doi: 10.1093/femsyr/foy048

Durán RV, Oppliger W, Robitaille AM, Heiserich L, Skendaj R, Gottlieb E, Hall MN (2012) Glutaminolysis activates Rag-mTORC1 signaling. Mol Cell 47: 349–358. doi: 10.1016/j.molcel.2012.05.043

Fernandez J, Yang KT, Cornwell KM, Wright JD, Wilson RA (2013) Growth in rice cells requires de novo purine biosynthesis by the blast fungus Magnaporthe oryzae. Sci Rep 3: 2398. doi: 10.1038/srep02398

Fernandez J, Marroquin-Guzman M, Wilson RA (2014) Evidence for a transketolase-mediated metabolic checkpoint governing biotrophic growth in rice cells by the blast fungus *Magnaporthe oryzae*. PLoS Pathog 10: e1004354. doi: 10.1371/journal.ppat.1004354

Fernandez J, Orth K (2018) Rise of a Cereal Killer: The Biology of *Magnaporthe oryzae* Biotrophic Growth. Trends Microbiol 26: 582–597. doi: 10.1016/j.tim.2017.12.007

Giraldo MC, Dagdas YF, Gupta YK, Mentlak TA, Yi M, Martinez-Rocha AL, Saitoh H, Terauchi R, Talbot NJ, Valent B (2013) Two distinct secretion systems facilitate tissue invasion by the rice blast fungus *Magnaporthe oryzae*. Nat Commun 4: 1996. doi: 10.1038/ncomms2996

Huang K, Czymmek KJ, Caplan JL, Sweigard JA, Donofrio NM (2011) HYR1-mediated detoxification of reactive oxygen species is required for full virulence in the rice blast fungus. PLoS Pathog 7: e1001335. doi: 10.1371/journal.ppat.1001335

Kankanala P, Czymmek K, Valent B (2007) Roles for rice membrane dynamics and plasmodesmata during biotrophic invasion by the blast fungus. Plant Cell 19: 706–724. doi: 10.1105/tpc.106.046300

Khang CH, Berruyer R, Giraldo MC, Kankanala P, Park SY, Czymmek K, Kang S, Valent B (2010) Translocation of *Magnaporthe oryzae* effectors into rice cells and their subsequent cell-to-cell movement. Plant Cell 22: 1388–1403. doi: 10.1105/tpc.109.069666

Kim B, Lee Y, Choi H, Huh WK (2021) The trehalose-6-phosphate phosphatase Tps2 regulates *ATG8* transcription and autophagy in *Saccharomyces cerevisiae*. Autophagy 17: 1013–1027. doi: 10.1080/15548627.2020.1746592

Klionsky DJ et al (2020) Guidelines for the use and interpretation of assays for monitoring autophagy. Autophagy 8: 445–544. doi: 10.4161/auto.19496

Li G, Marroquin-Guzman M, Wilson RA (2015) Chromatin Immunoprecipitation (ChIP) Assay for Detecting Direct and Indirect Protein – DNA Interactions in *Magnaporthe oryzae*. Bio-protocol 5: e1643.

Li G, Qi X, Sun G, Rocha RO, Segal LM, Downey KS, Wright JD, Wilson RA (2020) Terminating rice innate immunity induction requires a network of antagonistic and redox-responsive E3 ubiquitin ligases targeting a fungal sirtuin. New Phytol 226: 523–540. doi: 10.1111/nph.16365

Liao Y, Smyth GK, Shi W (2013) The Subread aligner: fast, accurate and scalable read mapping by seed-and-vote. Nucleic Acids Res 41: e108. doi: 10.1093/nar/gkt214.

Liao Y, Smyth GK, Shi W (2014) featureCounts: an efficient general purpose program for assigning sequence reads to genomic features. Bioinformatics 30:923–30. doi: 10.1093/bioinformatics/btt656

Loewith R, Hall MN (2011) Target of rapamycin (TOR) in nutrient signaling and growth control. Genetics 189: 1177–1201. doi: 10.1534/genetics.111.133363

Love MI, Huber W, Anders S (2014) Moderated estimation of fold change and dispersion for RNA-seq data with DESeq2. Genome Biol 15: 550. doi: 10.1186/s13059-014-0550-8

Marroquin-Guzman M, Hartline D, Wright JD, Elowsky C, Bourret TJ, Wilson RA (2017) The *Magnaporthe oryzae* nitrooxidative stress response suppresses rice innate immunity during blast disease. Nat Microbiol 2: 17054. doi: 10.1038/nmicrobiol.2017.54

Marroquin-Guzman M, Krotz J, Appeah H, Wilson RA (2018) Metabolic constraints on *Magnaporthe* biotrophy: loss of *de novo* asparagine biosynthesis aborts invasive hyphal growth in the first infected rice cell. Microbiology 164: 1541–1546. doi: 10.1099/mic.0.000713

Martin M (2011) Cutadapt removes adapter sequences from high-throughput sequencing reads. EMBnet.journal 17: 10–12. DOI: https://doi.org/10.14806/ej.17.1.200

May AI, Prescott M, Ohsumi Y (2021) Autophagy facilitates adaptation of budding yeast to respiratory growth by recycling serine for one-carbon metabolism. Nat Commun 11: 5052. doi: 10.1038/s41467-020-18805-x

Mittal N, Guimaraes JC, Gross T, Schmidt A, Vina-Vilaseca A, Nedialkova DD, Aeschimann F, Leidel SA, Spang A, Zavolan M (2017) The Gcn4 transcription factor reduces protein synthesis capacity and extends yeast lifespan. Nat Commun 8:457. doi: 10.1038/s41467-017-00539-y

Mora J (1990) Glutamine metabolism and cycling in *Neurospora crassa*. Microbiol Rev 54: 293–304. doi: 10.1128/mr.54.3.293-304.1990

Natarajan K, Meyer MR, Jackson BM, Slade D, Roberts C, Hinnebusch AG, Marton MJ (2001) Transcriptional profiling shows that Gcn4p is a master regulator of gene expression during amino acid starvation in yeast. Mol Cell Biol 21: 4347–4368. doi: 10.1128/MCB.21.13.4347-4368.2001

Pedruzzi I, Dubouloz F, Cameroni E, Wanke V, Roosen J, Winderickx J, De Virgilio C (2003) TOR and PKA signaling pathways converge on the protein kinase Rim15 to control entry into G0. Mol Cell 12:1607–1613. doi: 10.1016/s1097-2765(03)00485-4

Rocha RO, Wilson RA (2020) *Magnaporthe oryzae* nucleoside diphosphate kinase is required for metabolic homeostasis and redox-mediated host innate immunity suppression. Mol Microbiol 114: 789–807. doi: 10.1111/mmi.14580

Sakulkoo W, Osés-Ruiz M, Oliveira Garcia E, Soanes DM, Littlejohn GR, Hacker C, Correia A, Valent B, Talbot NJ (2018) A single fungal MAP kinase controls plant cell-to-cell invasion by the rice blast fungus. Science 359: 1399–1403. doi: 10.1126/science.aaq0892

Saunders DG, Aves SJ, Talbot NJ (2010) Cell cycle-mediated regulation of plant infection by the rice blast fungus. Plant Cell 22: 497–507. doi: 10.1105/tpc.109.072447

Srinivasan R, Walvekar AS, Rashida Z, Seshasayee A, Laxman S (2020) Genome-scale reconstruction of Gcn4/ATF4 networks driving a growth program. PLoS Genet 16: e1009252. doi: 10.1371/journal.pgen.1009252

Sun G, Elowsky C, Li G, Wilson RA (2018) TOR-autophagy branch signaling via Imp1 dictates plant-microbe biotrophic interface longevity. PLoS Genet 14: e1007814. doi: 10.1371/journal.pgen.1007814

Swinnen E, Wanke V, Roosen J, Smets B, Dubouloz F, Pedruzzi I, Cameroni E, De Virgilio C, Winderickx J (2006) Rim15 and the crossroads of nutrient signalling pathways in *Saccharomyces cerevisiae*. Cell Div 1: 3. doi: 10.1186/1747-1028-1-3

Tan HWS, Sim AYL, Long YC (2017) Glutamine metabolism regulates autophagy-dependent mTORC1 reactivation during amino acid starvation. Nat Commun 8: 338. doi: 10.1038/s41467-017-00369-y

Taus T, Köcher T, Pichler P, Paschke C, Schmidt A, Henrich C, Mechtler K (2011) Universal and confident phosphorylation site localization using phosphoRS. J Proteome Res 10: 5354–5362. doi: 10.1021/pr200611n

Tallóczy Z, Jiang W, Virgin HW 4th, Leib DA, Scheuner D, Kaufman RJ, Eskelinen EL, Levine B (2002) Regulation of starvation- and virus-induced autophagy by the eIF2*α* kinase signaling pathway. Proc Natl Acad Sci U S A 99: 190–195. doi: 10.1073/pnas.012485299

Valent B, Farrall L, Chumley FG (1991) *Magnaporthe grisea* genes for pathogenicity and virulence identified through a series of backcrosses. Genetics 127: 87–101. doi: 10.1093/genetics/127.1.87

Wanke V, Cameroni E, Uotila A, Piccolis M, Urban J, Loewith R, De Virgilio C (2008) Caffeine extends yeast lifespan by targeting TORC1. Mol Microbiol 69: 277–285. doi: 10.1111/j.1365-2958.2008.06292.x

Wilson RA, Talbot NJ (2009) Under pressure: investigating the biology of plant infection by Magnaporthe oryzae. Nat Rev Microbiol 7: 185–95. doi: 10.1038/nrmicro2032

Wilson RA, Fernandez J, Quispe CF, Gradnigo J, Seng A, Moriyama E, Wright JD (2012) Towards defining nutrient conditions encountered by the rice blast fungus during host infection. PLoS One 7: e47392. doi: 10.1371/journal.pone.0047392

Wilson RA (2021) Magnaporthe oryzae. Trends Microbiol 29: 663–664. doi: 10.1016/j.tim.2021.03.019

Yang Z, Geng J, Yen WL, Wang K, Klionsky DJ (2010) Positive or negative roles of different cyclin-dependent kinase Pho85-cyclin complexes orchestrate induction of autophagy in *Saccharomyces cerevisiae*. Mol Cell 38: 250–264. doi: 10.1016/j.molcel.2010.02.033

Yi M, Valent B (2013) Communication between filamentous pathogens and plants at the biotrophic interface. Annu Rev Phytopathol 51: 587–611. doi: 10.1146/annurev-phyto-081211-172916

Yorimitsu T, Zaman S, Broach JR, Klionsky DJ (2007) Protein kinase A and Sch9 cooperatively regulate induction of autophagy in *Saccharomyces cerevisiae*. Mol Biol Cell 18: 4180–4189. doi: 10.1091/mbc.e07-05-0485

